# An investigation of the sex-specific genetic architecture of fitness in *Drosophila melanogaster*

**DOI:** 10.1101/2022.08.04.502812

**Authors:** Amardeep Singh, Asad Hasan, Aneil F. Agrawal

## Abstract

In dioecious populations, the sexes employ divergent reproductive strategies to maximize fitness and, as a result, genetic variants can affect fitness differently in males and females. Moreover, recent studies have highlighted an important role of the mating environment in shaping the strength and direction of sex-specific selection. Here, we measure adult fitness for each sex of 357 lines from the *Drosophila* Synthetic Population Resource (DSPR) in two different mating environments. We analyze the data using three different approaches to gain insight into the sex-specific genetic architecture for fitness: classical quantitative genetics, genomic associations, and a mutational burden approach. The quantitative genetics analysis finds that, on average segregating genetic variation in this population has concordant fitness effects both across the sexes and across mating environments. We do not find specific genomic regions with strong associations with either sexually antagonistic (SA) or sexually concordant (SC) fitness effects, yet there is modest evidence of an excess of genomic regions with weak associations, both with SA and SC fitness effects. Our examination of mutational burden indicates stronger selection against indels and loss-of-function variants in females than males.

## Introduction

The genetic architecture of fitness variation (e.g., the number and identity of loci, and their effect sizes in different contexts) remains poorly understood. Given that any locus in the genome has the potential to contribute to fitness, the genetic architecture of fitness is likely highly polygenic. However, this does not preclude the possibility that some variants have reasonably large effects. Investigating the genetic architecture of fitness may provide useful clues as to the forces responsible for the maintenance of segregating fitness variants (e.g., mutation-selection balance or various forms of balancing selection; Mackay 2001; Barton and Keightley 2002; Frankham 2009)

In dioecious populations, the genetic architecture of phenotypic traits, including fitness may differ substantially between the sexes. Sex differences in the genetic architecture of fitness depend on the sex-specific effect size and direction of segregating variants. For example, some variants may have sex-limited effects because they are sex-linked or contribute to the expression of sex-limited traits (e.g., seminal fluid proteins). Such variants can be important contributors to the variance in fitness in one sex but may have little or no influence on fitness in the other. Variants with purely sex-limited fitness effects may be relatively rare. Instead, alleles may often contribute to variance in fitness in both sexes. The evolutionary fate of such variants will depend on the strength and direction of selection they experience when expressed in each sex. In some cases, alleles at a locus may have opposing, sexually antagonistic, effects on fitness resulting in so-called ‘intra-locus sexual conflict’ (reviewed in Bonduriansky and Chenoweth 2009). Under specific conditions, such variants can be maintained by balancing selection, particularly when the effect sizes of variants are large (Kidwell et al. 1977; Connallon and Clark 2012). However, most genes are thought to serve similar functions in both sexes and disruptions to those functions are likely deleterious for both sexes (Rowe and Houle 1996; Whitlock and Agrawal 2009). While sexually concordant selection should remove genetic variation at such genes (Long et al. 2012; Connallon and Clark 2013), this class of loci may serve as a large mutational target and thus contribute substantially to the standing genetic variation in fitness via mutation-selection balance.

The primary objective of this study is to better understand fitness variation within and across the sexes. We do so by measuring the *f*adult fitness*f* (i.e., fitness conditional on reaching the adult stage) of outbred *Drosophila melanogaster*, constructed from 357 recombinant inbred line (RIL) haplotypes from the *Drosophila* Synthetic Population Resource (DSPR; King et al. 2012). Ideally, one would catalog every variant and measure its fitness effect in each sex, but that ideal is not logistically feasible. Instead, we pursue three different approaches that each offer a different lens through which to view the relationship of the genetic architecture for fitness across the sexes.

First, we used a classic quantitative genetics approach to provide a summary of how genetic architecture of fitness is related between the sexes. This approach has been used in a number of laboratory (e.g., Vieira et al. 2000; Wayne et al. 2001; Pischedda and Chippindale 2006; Long et al. 2009; Collet et al. 2016) and wild (e.g., Kruuk et al. 2000; Merilä and Sheldon 2000; McCleery et al. 2004; Foerster et al. 2007) populations and is useful for asking how much additive genetic variation in sex-specific fitness exists in a population. In addition, we used this approach to estimate the intersexual genetic correlation for fitness (*r*_*mf*_), which serves as a useful indicator of the overall concordance or antagonism of segregating genetic variation in a population. This correlation can be used to quantify the extent to which improvement in the fitness of one sex will help or hinder the improvement of the other, at least in the short term. Empirical estimates of *r*_*mf*_ have been mixed: although a survey of animal species found that, on average, estimates of *r*_*mf*_ were small but positive (Poissant et al. 2010), several studies have reported negative estimates of *r*_*mf*_ of substantial magnitude (e.g., Chippindale et al. 2001; Fedorka and Mousseau 2004; Brommer et al. 2007; Punzalan et al. 2014). The power and limitation of the intersexual correlation is that it is a summary statistic that is predictive of short-term evolution that neither requires nor reveals information about individual allelic effects. Indeed, the underlying ‘genetic details’ matter little with respect to short-term evolution but can substantially affect evolutionary outcomes on longer time-scales.

Next, we used genomic association mapping techniques in an attempt to gain resolution on specific loci underlying sex-specific fitness. Association mapping provides a route to identifying individual allelic effects and, in principle, can be used to find the number and magnitude of mutations with sexually concordant, antagonistic, and sex-limited effects. Recognizing that the genetic basis of fitness is likely to be highly polygenic, we are unlikely to have the statistical power to identify a multitude of small effect variants. However, we were particularly interested in identifying loci contributing to sexual antagonism and such variants maintained by balancing selection are expected to be of larger effects (Kidwell et al. 1977; Connallon and Clark 2012), possibly detectable via genome association mapping. This approach has recently proven fruitful in identifying candidate sexually antagonistic loci in this species (Ruzicka et al. 2019).

Though association mapping is severely limited in its ability to identify small effect variants, other approaches can provide insights. When sequence and fitness data are both available for a collection of genotypes, then one approach is to quantify the aggregate effect of specified classes that are suspected to affect fitness *a priori* (Yang et al. 2017; Brown and Kelly 2019). For example, one can use this approach to ask how the total number of loss-of-function variants (LoFs) in a genome affects fitness, which is equivalent to quantifying the average strength of selection against segregating LoFs. We are particularly interested in comparing the selection strength in males versus females. Strong competition for access to females has been postulated to increase the strength of selection on males relative to females on sexually concordant variants (reviewed Whitlock and Agrawal 2009), potentially increasing population mean fitness by reducing mutation load (Manning 1984; Agrawal 2001; Siller 2001) and accelerating adaptation (Lorch et al. 2003). Experimental studies using phenotypic marker mutations (e.g., Whitlock and Bourguet 2000; Pischedda and Chippindale 2005; Stewart et al. 2005; Sharp and Agrawal 2008; Hollis et al. 2009) and mutation accumulation lines (e.g., Enders and Nunney 2010; Mallet et al. 2011; Mcguigan et al. 2011; Sharp and Agrawal 2013; Grieshop et al. 2016) tend to support the notion of stronger selection in males than females, though there are potential concerns with these approaches (see Discussion).

In addition to examining sex differences in the genetic architecture of fitness, the second goal of this study was to investigate how the genetic architecture of fitness of each sex, as well as the relationship of these architectures across the sexes, was affected by the ecological context in which sexual interactions occur. Here, we estimated adult fitness of males and females in two mating environments: traditional fly culturing vials and larger cages that represent lower density and more structurally complex environments. Previous work has shown males and females interact differently in environments similar to the two used here (Yun et al. 2017; MacPherson et al. 2018). Male harm and sexual conflict are strong features of sexual interactions in standard fly vials but not in the more complex cages; these differences affect the rates of nonsexual adaptation and purging (Singh et al. 2017; Yun et al. 2018).

Our quantitative genetic analysis recovers appreciable levels of additive genetic variation in adult fitness for both male and female fitness and finds that the fitness consequences of this genetic variation tends to be concordant across the sexes as well as across environments. Our association mapping analysis did not reveal any statistically convincing associations between fitness and any specific genomic region. However, there is some evidence that both sexually concordant and sexually antagonistic regions exist. Our mutation burden analysis recovers a significant negative association between the burden of putatively deleterious variants and female, but not male, fitness in both mating environments.

## Methods

### DSPR Inbred Genotypes

Recombinant inbred line (RIL) genotypes used for measuring sex-specific outbred fitness were obtained from the *Drosophila* Synthetic Population Resource (DSPR). The creation of these lines has been described in detail elsewhere (King et al. 2012a). Briefly, eight parental genotypes from geographically disparate regions were crossed in a round robin design to generate an outbred population that was allowed to recombine for 50 generations. Inbred genotypes were generated from this outbred, synthetic population following 25 generations of fullsib mating. Following low-resolution RADseq genotyping of inbred genotypes, a hidden Markov model (HMM) was implemented to determine the probable parental ancestry at regularly spaced 10kb intervals (or haplotype blocks) along the genome of each line. The parental ancestry of the vast majority of 10kb blocks in most lines can be assigned to a parental source with high certainty. Here, we estimated sex-specific fitness of 357 lines. Although the majority of lines were assayed for fitness in both sexes and both environments, this was not true of all lines for logistical reasons (see Table S1 for number of lines assayed per sex and environment).

### Measuring Adult Fitness of DSPR Genotypes

Our goal was to obtain a reasonably comprehensive measure of “adult fitness” (i.e., fitness conditional on reaching the adult stage) with respect to the contexts we use for these assays. (See Fig. S1 for a schematic summary of the assay procedure.) To measure sex-specific outbred adult fitness of focal genotypes we first crossed females of each focal genotype to males from a common outbred population that was generated by mixing all DSPR founder genotypes. Focal males and females were then assayed in separate competitive fitness assays. Specifically, for each outbred genotype, we combined four focal males/females with twelve competitors of the same sex from an outbred laboratory population homozygous for a dominant *bw*^*D*^ marker. Focal and competitor genotypes competed for access to sixteen unmarked individuals of the opposite sex that came from an outbred laboratory population. For logistical reasons the source population for these individuals differed between male and female fitness assays: in male fitness assays we used females from an outbred laboratory population originally captured from a West African population (*Dahomey* stock) while in the female fitness assays we used males from an outbred stock generated by mixing DSPR founder genotypes.

All flies were collected at approximately the same age (∼2-3 days post eclosion) and held separately by sex in vials sprinkled with live yeast in groups of sixteen for 24 hours prior to the start of fitness assays. All flies for a replicate were then combined and allowed to interact for 6 days in either a standard *Drosophila* culture vials (28 × 95 mm, approximately 60 mL) that contained 7ml of food sprinkled with live yeast or a larger, complex mating cages. The cages were polypropylene ULINE^®^ deli containers (114.3 × 76.2 mm, approximately 500 mL) with two food cups containing 7mL of food each, sprinkled with live yeast. Replicates were maintained in a 12L:12D light cycle at 25°C. Fresh media, sprinkled with live yeast was provided on day one and three (fresh media was not provided on day three in cage assays due to logistical reasons). On day six, all females were removed and, to reduce subsequent larval densities, divided approximately evenly into two vials sprinkled with live yeast and allowed to oviposit for ∼24 hours. Starting ten days later, and repeating for a total of three days, offspring from the two oviposition vials were collected, pooled and frozen daily. The vast majority of offspring emerged on the first two days of collection. Following collection, we randomly sampled at least 100 offspring and calculated the proportion of total offspring with wildtype (red) eye colour in this sample. We have five replicate assays for most combinations of line, sex, and environment. Due to human error, some lines have more than five replicates while others have fewer. Fitness assays were conducted in a blocked design.

Though we consider these measures to be a reasonable representation of *f*adult fitness*f*, they are indirectly affected by juvenile survivorship because we assess the fitness of adult males and adult females by measuring the number of offspring that survive to the adult stage. However, we expect variation in juvenile survival to make only a small contribution to the genetic variance in our “adult fitness” measures for several reasons. First, we attempted to keep offspring density from being high so that survivorship should be high. Second, the contribution of the genetic variance in offspring survivorship to the variance in focal adult fitness will be diluted by 50% because offspring only have half the genes from the focal parent. Third, previous studies have indicated that genetic variation for juvenile survivorship is small, especially compared to that of adult fitness components (e.g., Leips and Mackay 2000; Sharp and Agrawal 2018). Though we expect the influence of juvenile fitness effects to be small, to the extent they contribute to the genetic variance, they will upwardly bias estimates of the intersexual genetic correlation (*r*_*mf*_) because both surviving sons and daughters contribute to fitness measures of each sex.

### Quantitative Genetic Analysis of Fitness in the DSPR

To estimate the sex-specific heritability in fitness in each mating environment, we fit separate mixed models for each mating environment to estimate the sex-specific additive genetic and residual variances. We modeled the proportion of offspring descended from focal (rather than competitor) flies as a binomial variable using *MCMCglmm* (Hadfield 2010). Sex was included as a fixed effect whereas block and RIL genotype were included as random effects. The model assumed the two sexes had separate and independent variances due to block (as the sexes were assayed in separate blocks with different competitors). On the logit scale, the model estimates RIL effects via a bivariate Gaussian distribution *N*(0, ***G***) where

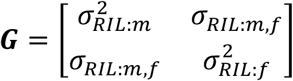

is a genetic covariance matrix. The diagonal elements are the variances among RILs for male and female fitness, which are proportional to the additive genetic variances; the off-diagonal is the among-RIL intersexual covariance, which is proportional to the additive genetic covariance. (Our estimated matrix is proportional, rather than equal, to the traditional ***G*** matrix because our unit of observation is the fitness of groups of four focal individuals sharing a RIL haploid genome rather than a single individual; see Supplemental Material for derivation). *MCMCglmm* allows for additive overdispersion via inclusion of a random effect (on the logit scale) for each observation; we assumed the sexes had independent variances in this regard. Models were fit using extended parameter priors (e.g., *V*=*diag*(2), *nu*=1, *alpha*.*mu* = c(0, 0), *alpha*.*V* = 1000\**diag*(2)), which are recommended when genetic variances are small (de Villemereuil 2018). (As a simple check, we permuted the replicate fitness values with respect to RIL identity within each block for each sex and ran the same analysis; inferred genetic variances were an order of magnitude smaller.) Because the values returned from *MCMCglmm* are on the link scale, we used *QGglmm* (de Villemereuil *et al* 2016) to obtain estimates on the observed scale.

Posterior distributions for parameters of interest were made from the MCMC chain of model terms, following conversion to the observed scale and any required subsequent calculation (e.g., as for heritability). Because we measure fitness from replicates containing four individuals, the additive genetic variance for sex *j* is 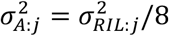 where 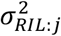 is the variance (on the observed scale) across all replicates associated with RIL identity (see Supplementary Material). Heritability in sex *j* is given by

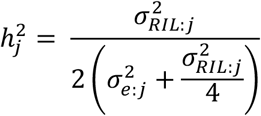

where 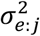 is the non-genetic variance across all replicates for sex *j*. Evolvability for sex *j* was calculated as 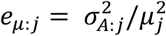 where *μ*_*j*_ is the mean adult fitness measure of an individual of sex *j*. The intersexual genetic correlation for fitness (*r*_*mf*_) is calculated as

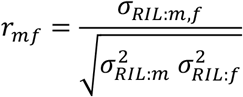

Reported point estimates are means of the posterior and 95% credibility intervals (CI) were calculated using the *HPDInterval* function in *MCMCglmm*.

Though our estimates of *r*_*mf*_ are intended to represent the intersexual genetic correlation with respect to “adult” fitness, they are indirectly affected by juvenile survivorship because we assess the fitness of adult males and adult females by measuring the number of offspring that survive to the adult stage. Genetic variation for offspring survivorship will upwardly bias our estimate of the intersexual correlation for adult fitness. However, we expect this effect to be small for several reasons. First, we attempted to keep offspring density from being high so that survivorship should be high. Second, the contribution of the genetic variance in offspring survivorship to the variance in focal adult fitness will be diluted by 50% because offspring only have half the genes from the focal parent. Third, previous studies have indicated that genetic variation for juvenile survivorship is small, especially compared to that of adult fitness components (e.g., Leips and Mackay 2000; Sharp and Agrawal 2018).

As a separate question, we were interested in the relationship between the genetic architectures of a given sex across the two mating environments. Using the fitness data for a single sex from each of the two environments, we employed a similar approach to estimate the within-sex cross-environment genetic correlation for fitness (i.e., *r*_*env,m*_ and *r*_*env,f*_ for males and females, respectively).

### Association Mapping of Fitness and Indices of Sexual Antagonism and Concordance

We briefly outline methods for genome association mapping and provide details below. To identify genome associations with sex-specific fitness as well as loci underpinning sexually antagonistic and concordant variation, we first identified univariate traits for fitness and indices of sexual antagonism and concordance. Next, we defined a statistical model for mapping genotypic variation to trait variation and a significance threshold was established to identify loci putatively associated with fitness or axes of antagonism/concordance.

To map phenotypic variation in male and female fitness we calculated the mean relative fitness, separately for each sex, of each RIL. To associate loci with sexual antagonism and sexual concordance, we defined an ‘axis of antagonism’ (i.e., SA Axis) and an ‘axis of concordance’ (i.e., SC Axis) following the method by Berger et al. (2014), which we illustrate in Fig. S2. The *SC* value of a RIL is computed as 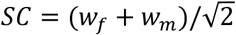 where *w*_*f*_ and *w*_*m*_ are the estimates of mean female and male relative fitness. This value represents the distance of a genotype from mean in the bivariate space of male and female relative fitnesses in the direction of equal male and female fitness effects (i.e., the 1 to 1 line). RILs with above (below) average fitness for both sexes are expected to have positive (negative) values of SC. The *SA* value of a RIL is calculated as 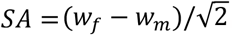. This value represents the distance of a genotype from mean in the direction of equal but opposite female and male fitness effects (i.e., the *w*_*f*_ = -*w*_*m*_ line). RILs with above (below) average values for female fitness but the opposite for male fitness are expected to have positive (negative) values of SA. Our primary interest was with respect to the SA index, but we independently mapped four traits—SA index and SC index, as well as male and female fitnesses—separately for each environment (i.e., 8 total traits).

A feature of the DSPR is that the mosaic haplotype structure of each RIL is known to a high degree of certainty in most cases. Specifically, at 10kb intervals along the genome of each RIL, the additive probability of each of the eight parental genotypes is known from a Hidden Markov Model (King et al. 2012a). To associate loci with each of the eight traits we regressed, at each 10kb haplotype block, the RIL trait values against the additive probability for each of the eight parental lines. Specifically, we fit the mixed model:

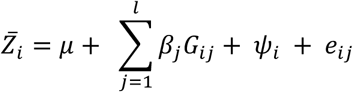

where 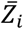 is the mean trait value of the *i*th RIL and *μ* is the grand trait mean across all sampled RILs, *G*_*ij*_ is fixed effect of the additive probability of the haplotype block of the *i*th RIL having descended from parental genotype *j, β*_*j*_ is the additive genetic effect of the *j*th parental genotype on the trait, *ψ*_*i*_ is the random effect of RIL genotype that is assumed to follow a Gaussian distribution such that *ψ*_*i*_ ∼ *σ*_*kl*_ ∼ *N*(0, ***K***) where ***K*** is the relatedness matrix of the DSPR lines (see below), *l* is the number of parental genotypes (max = 8), that contributed a cumulative additive probability of at least 5% at each haplotype block *i* across all RIL genotypes (see below) and *e*_*ij*_ specifies the residual variance of the *i*th RIL at the *j*th haplotype block and is assumed to follow a Gaussian distribution with a mean of zero and unknown variance. At each 10kb haplotype block *i*, we excluded any RIL genotypes whose most probable parental ancestor did not have a cumulative additive probability of at least 5% among all RIL genotypes. *P*-values were obtained from a likelihood ratio test against a null model where *β*_*j*_ = 0 for all *j*.

To estimate the genetic relatedness matrix of lines assayed, we calculated the average pairwise relatedness for all 357 inbred genotypes over all haplotype blocks. Specifically, for each pair of genotypes *i* and *j* we calculated the genetic relatedness (*κ*_*ij*_) as

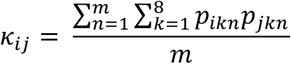

where *p*_*ikn*_ is the additive probability that haplotype block (i.e., a 10kb interval) *n* in genotype *i* originated from ancestral genotype *k* and *m* is the total number of haplotype blocks where genotypic information is present in both inbred genotypes.

As reported below, we were unable to find strong statistical support for individual haplotype blocks contributing to sexual concordance or antagonism. A plausible explanation is that effect sizes are low relative to environmental variation, so it becomes unlikely to find the very low *p*-values required when controlling for multiple testing. To explore the data, we shifted our focus to testing for an excess of haplotype blocks with *p*-values below a given critical value, *p*_*crit*_. In principle, the expected number of such blocks under the null hypothesis of no true genetic effects (i.e., the number of false positives, *x*) should be *E*[*x*] = *p*_*crit*_*n*_*total*_ where *n*_*total*_ is the number of blocks tested. If the tests were independent, it would be straightforward to calculate the full probability distribution of *x* under the null hypothesis and properly test whether the observed number of ‘significant’ blocks *n*_*p*≤*crit*_ is significantly larger than expected under the null hypothesis. There are two potential issues with this approach. First, it relies on the *p*-value from each individual test accurately reflecting the chance of a false positive under the null hypothesis (i.e., that all the assumptions of the mixed model are well met). Second, the tests are not independent due to linkage disequilibrium. Permutations offer a conceptually simple (though computationally intensive) solution to both issues. We permuted line identity (i.e., genome-wide genotype) with respect to the set of fitness values (i.e., the 8 traits) and reanalyzed the data. From each permutation, we recorded the *p*-value for each block (and for each trait) and then calculated an *x* value (i.e., number of ‘significant’ blocks) for a given statistical criteria (*p*_*crit*_ ∈ {0.001, 0.01, 0.05, 0.1}). To reduce computational time, we only tested every 7^th^ block; the same blocks were tested in the unpermuted and permuted data sets. (There is little additional information gained from testing all blocks because of the high linkage disequilibrium among neighboring blocks.) 2077 permutations were performed.

False discovery rates are an alternative approach to the multiple testing problem inherent in genomic analyses but are not necessarily a perfect solution. The reliability of these methods depend not only on the validity of the underlying *p*-values, but can depend on the independence of tests as well as the shape of the distribution of *p*-values. The *R* package *qvalue* (Storey et al. 2021) was used to convert *p-*values into *q*-values (i.e., FDR rate) based on a previously described method (Storey and Tibshirani 2003). We calculated *q*-values with the unpermuted data as well as each permuted data set. As we did with *p*-values, we compare the number of blocks with a *q*-value below a given threshold, *q*_*crit*_, to the distribution under the null hypothesis as determined from the permutations.

#### Gene Ontology (GO) Enrichment Analysis

We conducted a gene ontology (GO) enrichment analysis of genes found in *f*genomic clusters*f* that are associated with male and female fitness and the SA and SC axes. To help control for strong linkage between loci, we employed a clustering algorithm to define genomic clusters that are associated with each trait. For each chromosome arm, our clustering procedure began at the block furthest away from the centromere. We used a sliding window of 3.5 Mb to identify windows (or clusters) that had at least one haplotype block with *p* < 0.1. When a significant haplotype block was identified, we examined the immediately proceeding 3.5 Mb window. If this window contained any haplotype blocks that reached statistical significance, these blocks was subsumed into the previous cluster. This procedure was repeated until no haplotype blocks in the proceeding 3.5 Mb window reached statistical significance. Thus, a cluster was defined as a set of statistically significant haplotype blocks that are closer than 3.5 Mb to each other. After obtaining genomic clusters associated with each of the eight traits (see above), we used Ensembl’s *Variant Effect Predictor* (*VEP*; McLaren et al. 2010) to find genes that fall 5Kb upstream or downstream of the leading haplotype block in each cluster (i.e., the haplotype block with the lowest *p*-value). This analysis used a liberal *p* < 0.1 cut off for statistical significance of haplotype blocks but in many cases the leading block for a cluster would have a much lower *p-*value. Next, we used *GOrilla* (Eden et al. 2009) to test for an enrichment of GO terms in genes associated with each trait relative to a background set of all genes annotated in the *Drosophila* genome from our *VEP* analysis. *p*-values for GO terms were converted to *q-*values and a *q*-value threshold of 0.05 was used to assess statistical significance.

### Selection on Putatively Deleterious Variants in the DSPR lines

Our pipeline to identify putatively deleterious variants (i.e., single nucleotide polymorphisms [SNPs] and insertion/deletion [indel] variants) involved three steps. First, we imputed the underlying genomic sequence at sites segregating for SNPs and indels in the DSPR for all 357 focal genotypes assayed in this study. Following quality control steps, we attempted to enrich for sites likely to be deleterious by filtering out any variants segregating in either of two geographically disparate outbred populations because variants that are deleterious should be rare in nature. Finally, we mapped the variants to functional categories and retained only those variants falling within coding regions and whose functional consequence could reasonably be expected to be deleterious. The details of these steps are provided in the Supplementary Material.

Following these steps, we obtained a list of three classes of rare and putatively deleterious variants segregating in coding regions: loss-of-function (LoF) variants, indels, and variants resulting in radical amino acid changes. We scored each inbred genotype for the number of variants of each class that they carry. We termed the number of putatively deleterious variants of carried by a genotype to be that genotype’s “mutational burden”. Note that the estimated number of deleterious variants for each RIL will, on average, be slightly below the true number because a small fraction of genomic blocks are excluded from each RIL. The “uncounted” variants are a small source of measurement error in the estimate of burden but are not expected to create biases in the effects of interest (i.e., direction and magnitude of variants or sex differences in these effects). Two genotypes were clear outliers with respect to the number of SNPs causing radical amino acid changes (Figure S3) and as result these two genotypes were removed from further analysis.

To estimate the fitness effects of mutational burden, data from both sexes in both environments were analyzed in a single model. We used *MCMCglmm* (with family = “multinomial2”) to perform a mixed model analysis that included *n*_*LoF*_, *n*_*indel*_, *n*_*radical*_ (i.e., numbers of variants of each class) as well as fixed effects of sex, environment, sex-×-environment and the interaction of these with variant numbers. The random effects (and priors) were similar to the models described in the quantitative genetics section. Because both environments were included here, this model allowed for separate and independent ‘block’ and ‘unit’ (replicate) variances for each of the four sex-×-environment combinations. The model also allowed for RIL-associated variances for each sex-×-environment combination as well as all of their covariances. The expected fitness, on the observed scale, of a replicate of sex *i* in environment *j* with a given mutational burden can be calculated as:

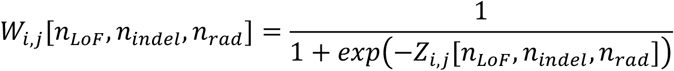

with

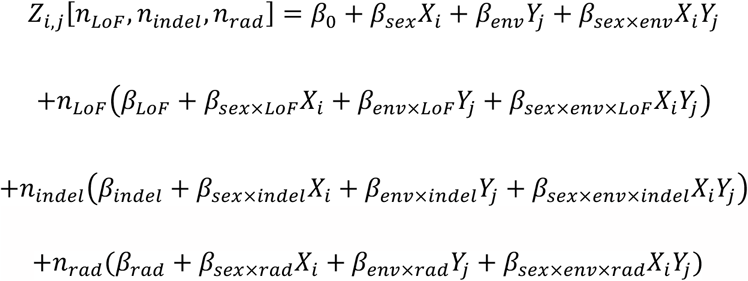

where the *β*’s are the estimated model coefficients (on the logit link scale) and *X*_*i*_ is an indicator variable for sex (*X*_*i*_ = 0 for females and 1 for males) and *Y*_*j*_ is an indicator variable for sex (*Y*_*j*_ = 0 for complex and 1 for simple). Selection against LoFs for sex *i* in environment *j* can be estimated as

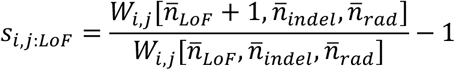

Selection for the other two variant classes can be estimated in the same fashion. Given our interests, for each variant class, we focus on three measures: (i) selection averaged over both sexes and environments: *s* = (*s*_*f,comp*_ + *s*_*f,simp*_ + *s*_*m,comp*_ + *s*_*m,simp*_)/4, (ii) sex differences in selection: Δ_*sex*_= *s*_*f*_ − *s*_*m*_ (this is averaging over environments, i.e., *s*_*f*_ = (*s*_*f,comp*_ + *s*_*f,simp*_)/2); and (iii) selective differences between environments Δ_*env*_= *s*_*comp*_ − *s*_*simp*_ (this is averaging over sexes, i.e., *s*_*comp*_ = (*s*_*f,comp*_ + *s*_*m,comp*_)/2). For each of the three variant classes, we estimated *s*, Δ_*sex*_ and Δ_*env*_ from each saved iteration in the MCMC chain to generate posterior distributions.

## Results

### Quantitative Genetic Analysis

We recover modest additive genetic variation 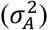 for adult fitness (measured as number of offspring produced) in both sexes when assayed in simple 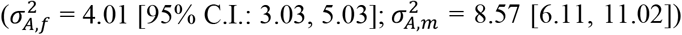 and complex 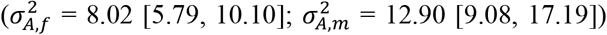 mating environments. We confirmed by visual inspection that the posterior distribution for each genetic variance component was unimodally distributed with similar mode and mean values. These genetic variances correspond to modest estimates of heritability (simple environment: 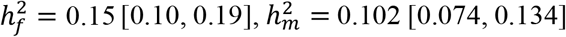; complex environment: 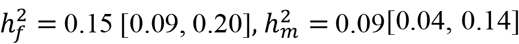 and evolvability (simple environment: *e*_*μ:f*_ = 0.05 [0.04, 0.06], *e*_*μ:m*_ = 0.06 [0.04, 0.08]; complex environment: *e*_*μ:f*_ = 0.08 [0.06, 0.10], *e*_*μ:m*_ = 0.08 [0.05, 0.11]).

The intersexual genetic correlation for fitness was positive and significantly different from zero in both mating environments (Figure 1; simple: *r*_*mf*_ = 0.31 [0.14, 0.48]; complex: *r*_*mf*_ = 0.21 [0.03, 0.38]). Though positive with credibility intervals that do not overlap zero, these correlations are far from unity, suggesting a fair degree of genetic independence between the sexes. The cross-environment correlation for fitness was also positive and substantially different from zero and unity in both sexes (Figure 2; *r*_*env,f*_ = 0.28 [0.12, 0.45], *r*_*env,m*_ = 0.40 [0.21, 0.57]).

**Figure 1:**
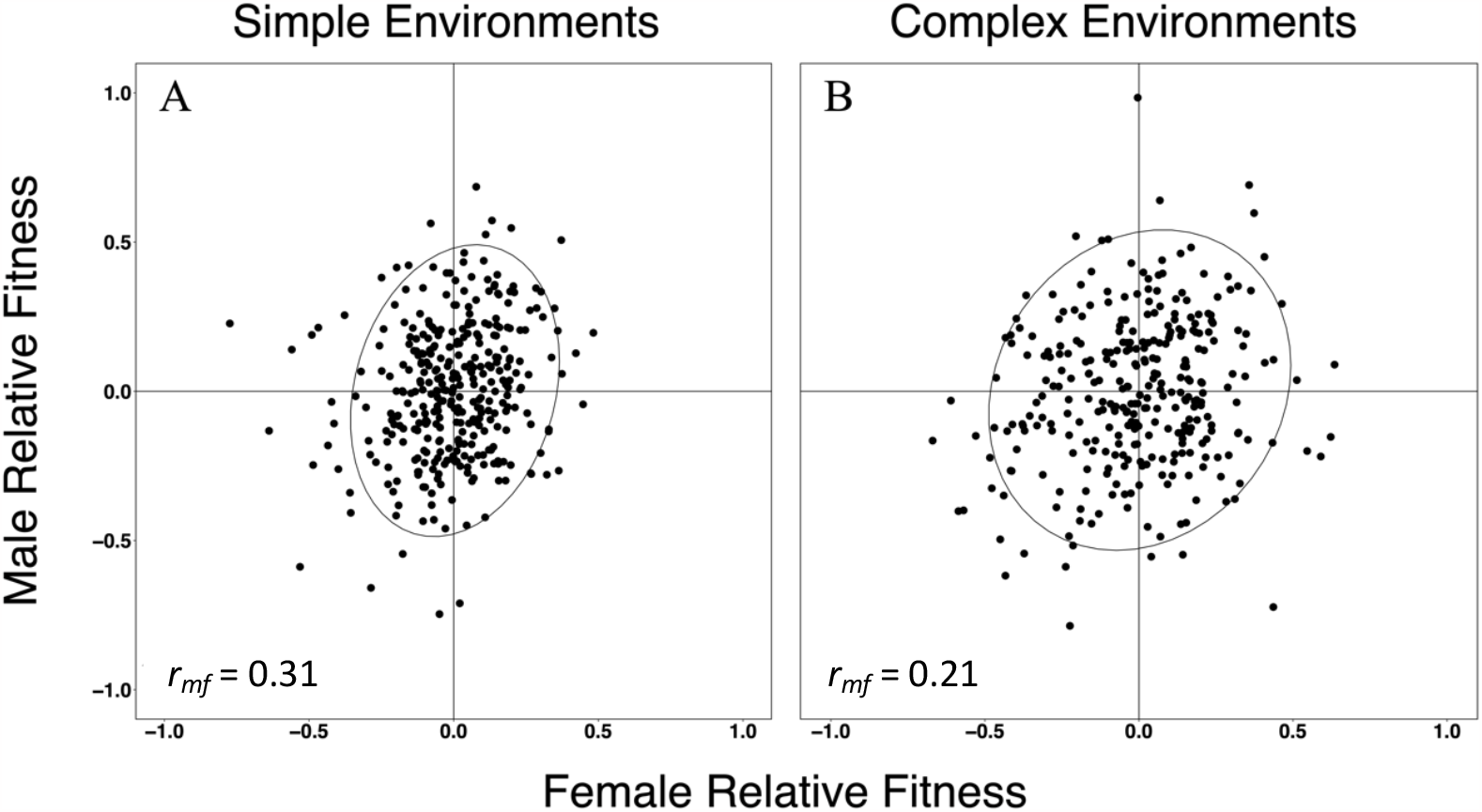
Male and female mean relative fitness of DSPR genotypes measured in **(A)** simple and **(B)** complex mating environments. For this figure (but not analysis), fitness measures are standardized to the within-block mean fitness for each sex and environment respectively and zero centered. Ellipses represent the 95% confidence ellipse for the distribution of sex-specific mean fitness estimates.

**Figure 2:**
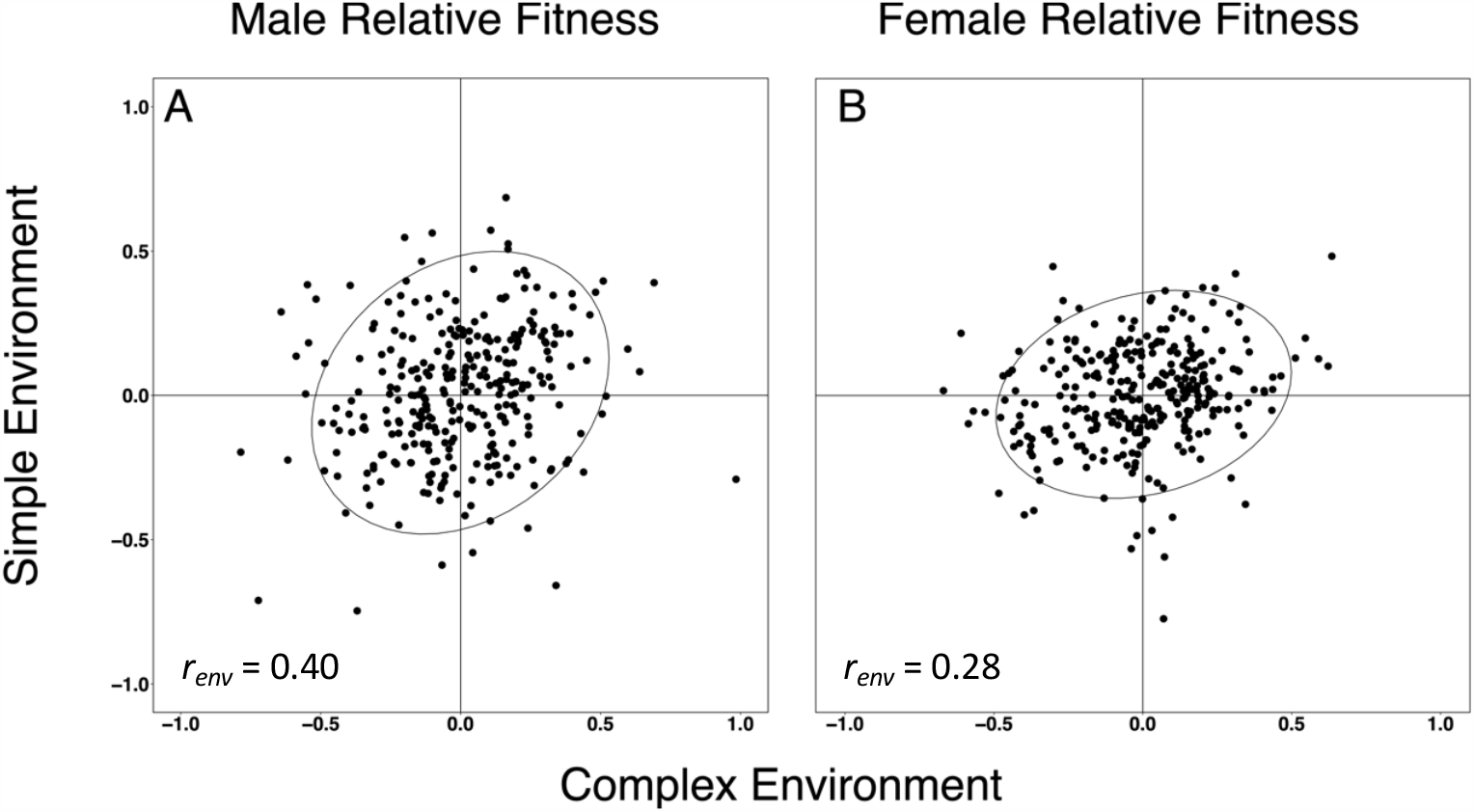
Mean relative fitness measured in simple and complex mating environments in **(A)** males and **(B)** Females. For this figure, fitness measures are standardized to the within-block mean fitness for each sex and environment respectively and zero centered. Ellipses represent the 95% confidence ellipse estimated from the sex-specific mean fitness estimates.

### Association Mapping of Sexually Antagonistic and Sexually Concordant Loci

We present the results for sexual antagonism (SA) and sexual concordance (SC) here and results for male and female fitness are given in the Supplemental Materials. Manhattan plots for SA and SC in the simple and complex environments are shown in Figure 3. Few haplotype blocks have strong statistical associations with SA or SC; thus, we have little confidence in specific regions affecting SA or SC. Consequently, we shifted our focus to exploring whether the there is any evidence at all of regions associated with SA or SC.

**Figure 3:**
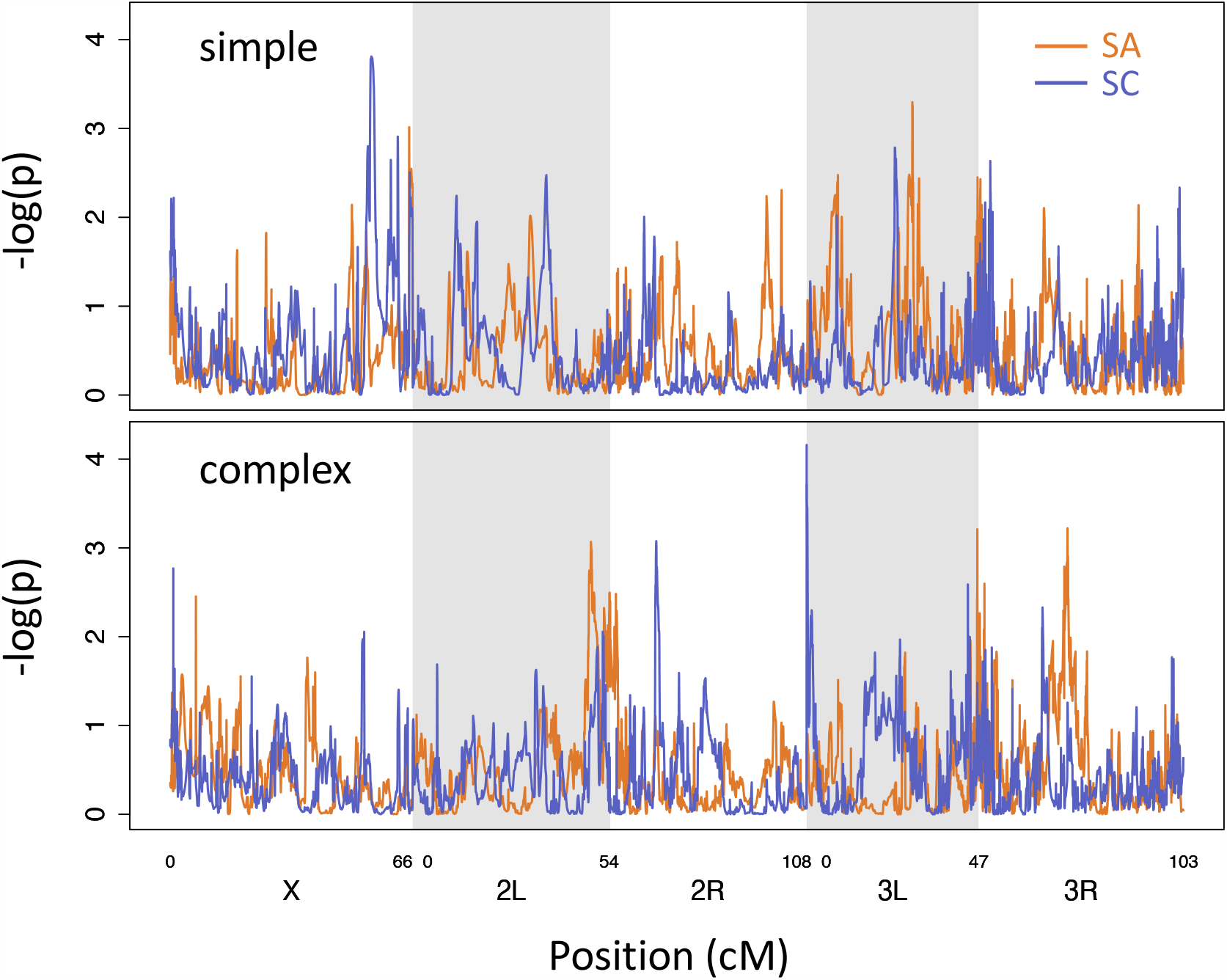
Manhattan plots. Strength of statistical support for associations between 10kb haplotype blocks along the genome with sexual antagonism (SA; orange) and sexual concordance (SC; purple) in DSPR lines when fitness was measured in the simple (top) or complex (bottom) mating environment.

As reported above, the the classical quantitative genetic analysis indicates the existence genetic variation for both sexes. Because SA and SC are simple linear transformations of female and male fitnesses, we can readily compute the additive genetic variances of the former from the latter (simple environment: 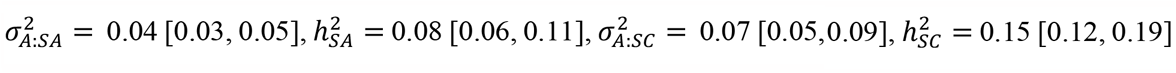; complex environment: 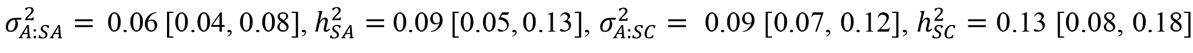; see Supplementary Material for details of calculations). Given the existence of this quantitative genetic variation for SA and SC, it is plausible that many blocks have modest effects that do not yield extreme *p*-values (e.g., *p* < 10^−5^). To explore this possibility, we counted the number haplotype blocks *n*_*p*≤*pcrit*_ that passed a given statistical requirement (e.g., *p*_*crit*_ *=* 0.001, *p*_*crit*_ *=* 0.05) and compared this number to the distribution under the null hypothesis of no genetic effects as generated by permutation.

Consider the results using *p*_*crit*_ *=* 0.05. We first ask if there is evidence of genetic associations of any sort, considering the tests for all four traits (i.e., SA and SC in both environments). We find 610 tests that pass the criterion of *p* ≤ 0.05 (“significant blocks”). In permutations, the average number of significant blocks is about half this number (353.3) and fewer than 1% of permutations result in 610 or more blocks (i.e., there are more associations than expected by chance). Examining SA and SC separately, there are 338 significant associations with SA considering both environments. In permuted data sets, the average number of SA blocks is 176.2 and fewer than 2% of permutations have 338 or more. There are 272 significant SC blocks across environments though ∼10% of permutations have as many or more. Examining SA and SC in each of the environments separately, there are more observed blocks than the corresponding average number observed for each of the four traits, but only for SA in the simple environment is the observed number in the upper 5% of the tail of its corresponding permutation distribution (Table 1).

**Table 1.**
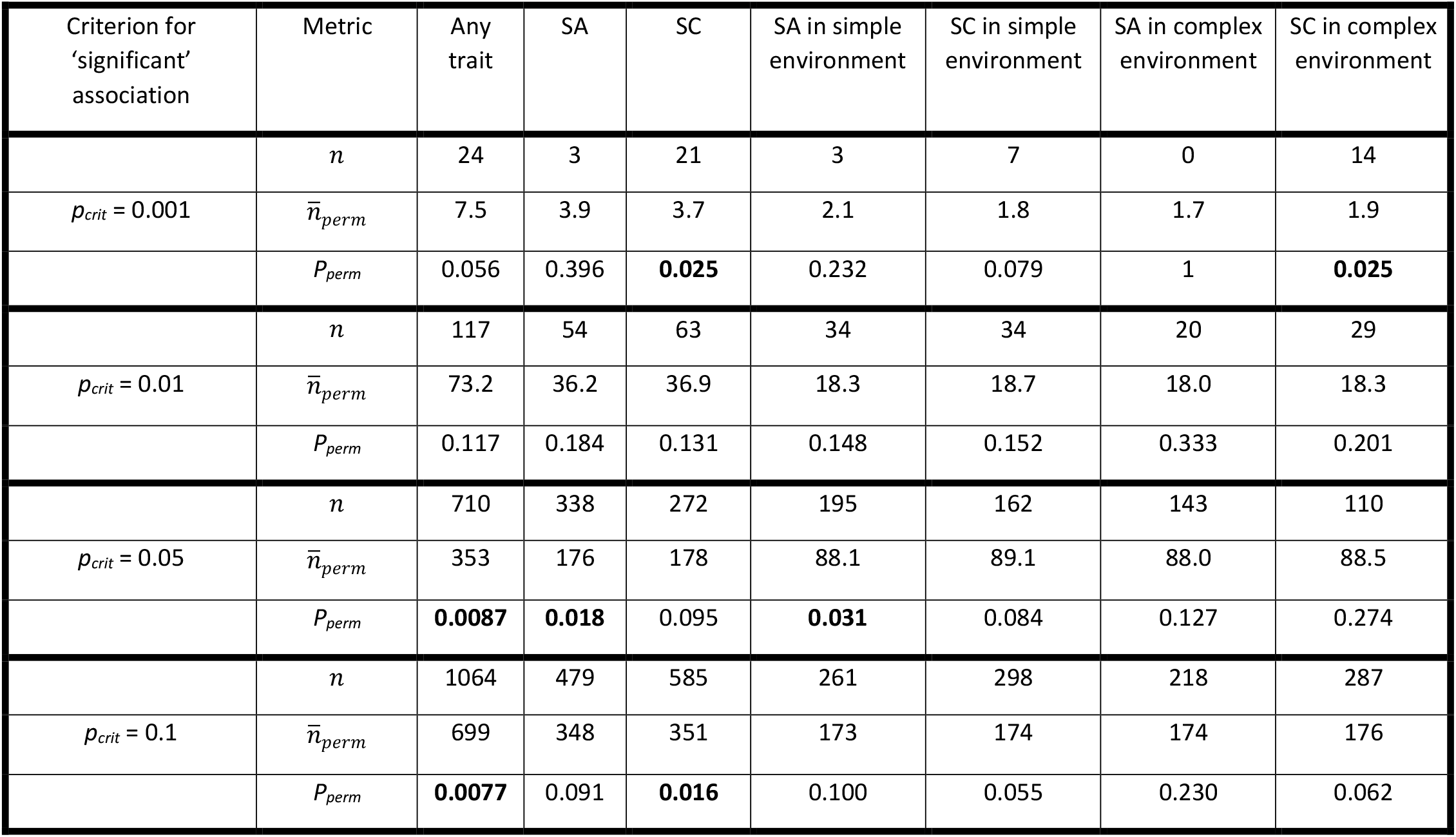
The number haplotype blocks passing a specified statistical criteria. For this analysis, only every 7^th^ haplotype block was tested (1682 blocks tested). *n* is the observed number in the real (i.e., unpermuted data). 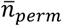 is the mean number of associations in permutated data sets. *P*_*perm*_ is the proportion of permutations with as many as or more than the observed number of associations (*n*); cases where the observed value is the upper 5% of the distribution (i.e., *P*_*perm*_ < 0.05) are shown in bold. Results using a *q-*value criterion are shown in the Supplementary Material, as are results for male and female fitness (Tables S2-4).

These results are sensitive to the statistical criterion (*p*_*crit*_) used to identify significant blocks (Table 1). In most cases, the observed number of blocks is greater than expected (i.e., the means of the permutation distributions) but whether the excess is substantially greater than expected depends on which trait is being considered and what statistical criterion is used. Whereas the results based on *p*_*crit*_ = 0.05 indicate an excess of associations for SA but not SC, with *p*_*crit*_ = 0.001 or *p*_*crit*_ = 0.1, there appears to be an excess of associations for SC but not SA. With *p*_*crit*_ = 0.01, neither type of association is in the upper 5% of the tail of the permutation distribution.

The use of permutations as a basis for a null distribution is important in evaluating the data from the DSPR. As an example, consider the associations tests for SA in the simple environment using *p*_*crit*_ = 0.05. 1682 blocks are tested. Under the null hypothesis of no true associations, we expect 1682*0.05 = 93.1 ‘significant’ blocks on average; the mean of the permutations is close to this value at 88.1. If the tests were independent, then the variance among permutations should be 1682*0.05*0.95 = 88.5; however, the observed variance—1952—is much larger. This higher variance is likely due to the non-independence of tests because of linkage disequilibrium, which can result in the false positive rate being much higher (or much lower) than expected under independence in any given instance (i.e., individual permutation or real data). The use of *q*-values does not solve this issue (Table S9). For example, applying the significance criterion of *q* ≤ 0.3, only 73% of permutations result in 0 ‘significant’ blocks and, of the remaining 27% of permutations, the average number of ‘significant’ blocks is 129 (range: 1-1675).

Following the procedure outlined in the Methods to generate lists of candidate regions, we conducted GO enrichment analyses (Tables S5-S12). Under the statistical criterion used, no GO terms were enriched among genes associated with the SA and SC in the simple environment. However, we find evidence of enrichment of GO terms among genes associated with SA and with SC in the complex environment. For SA, the enriched categories are biological processes that affect basic cell function as well as higher levels of regulation (e.g., post-translational regulation, epigenetic regulation; Table S7). For SC, a single molecular process was enriched: dipeptidyl-peptidase activity (Table S8). Though the terms enriched are not readily interpretable, the existence of enriched terms is another line of evidence that the association mapping results are more than mere noise.

### Mutational Burden

The assayed DSPR genotypes had on average 5.6 (range: 1-12) LOFs, 125.7 (75-172) indels in coding regions, and 274.8 (39-363) radical amino acid missense variants. We estimated the fitness effects of each of these three variant classes with the goal of addressing three questions: What is the average fitness effect per variant (*s*), averaging over both sexes and both environments? What is the difference in fitness effects between sexes (*Δ*_*sex*_ = *s*_*f*_ – *s*_*m*_)? What is the difference in fitness effects between environments (*Δ*_*env*_ = *s*_*comp*_ – *s*_*simp*_)?

For the two variant types with the strongest *a priori* expectation to be deleterious—LoFs and indels—the point estimates of average selection (*s*) are, counter to expectation, slightly positive but with 95% credible intervals broadly overlapping zero (Figure 4). However, for both LoFs and indels, there is evidence that deleterious selection is stronger in females than males (LoFs: *Δ*_*sex*_ = -0.021 [-0.038, -0.003]; indels: *Δ*_*sex*_ = -0.0019 [-0.0038, -0.0001]). In females, adult fitness is estimated to be reduced by ∼ 0.7% per LoF and ∼0.06% per indel; in males, the estimates are of the opposite sign and with substantially higher uncertainty. For these two variant types, there is no support for environmental differences (i.e., credibility intervals for *Δ*_*env*_ overlap zero). For radical amino acid missense variants, average selection is estimated to be positive rather than negative (*s* = 0.0008 [0.0001, 0.0014]). For this variant type, there is no support for sex differences (credibility intervals for *Δ*_*sex*_ overlap zero) but selection appears to differ between environments (*Δ*_*env*_ = -0.0015 [-0.0028, -0.0001]), with selection favouring such variants more strongly in the simple than complex environment.

**Figure 4:**
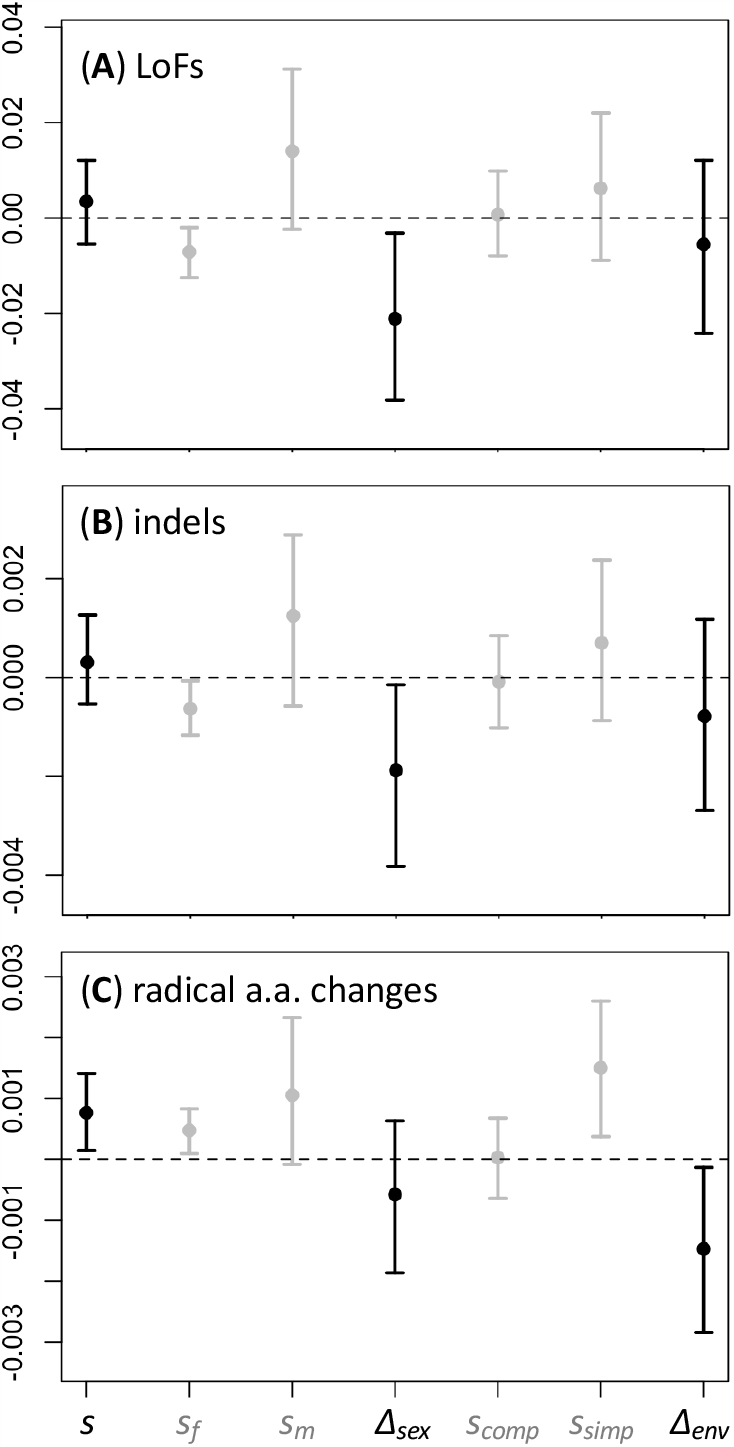
Fitness effects of putatively deleterious variants: (A) LoFs, (B) indels, (C) radical amino acid missense variants. *s* is the estimated selective effect per variant, averaging across sexes and environments (negative values indicate deleterious fitness effects); *s*_*f*_ (*s*_*m*_) is the selection per variant in females (males), averaging across environments; *Δ*_*sex*_ = *s*_*f*_ – *s*_*m*_ is the sex difference in selection; *s*_*comp*_ (*s*_*simp*_) is the selection per variant in the complex (simple) environment, averaging across sexes; *Δ*_*env*_ = *s*_*comp*_ – *s*_*simp*_ is the sex difference in selection.

## Discussion

The sexes represent different selective environments: surveys of phenotypic selection have found that shared traits experience persistent sex differences in both the strength (Singh and Punzalan 2018) and direction (Cox and Calsbeek 2009; De Lisle et al. 2018) of selection. Thus, there is considerable scope for segregating genetic to affect fitness of the two sexes differently. On the other hand, the two sexes share many similar requirements so selection on many variants will be similar. In this study, we used three approaches to explore different aspects of the sex-specific genetic architecture of fitness in a ‘synthetic’ laboratory population of *Drosophila melanogaster*.

The history of this synthetic population should be considered in interpreting our results. The RILs of the DSPR were established after first mixing eight inbred lines from different geographic regions and allowing for 50 generations of recombination (King et al. 2012a,b). During that period, some level adaptation to laboratory conditions is likely to have occurred. The selection regime that occurred during period was not identical to either of the environments used in this assay, though presumably it was more similar to those environments than to selection in many environments in nature where this species is found. Thus, it is difficult to say how well adapted the population was to the assay conditions. This issue is not unique to our study; most studies lack a quantitatively meaningful measure of how far a population is from maximal adaptation, though it is common to claim a lab population of invertebrates is “well-adapted” if it has been maintained in the lab for a few tens of generations.

This is an important issue because the genetic variation will be shaped by the history of selection. Selection is expected to drive variants that are deleterious in a given environment to low frequency. Thus, a population with a history of selection in a given environment may carry many fewer such variants at intermediate frequency than a population without that history of selection. This issue is not only relevant to the amount of genetic variance for fitness but also the intersexual genetic correlation. Populations experiencing a new environment may harbour numerous variants that are deleterious for both sexes. Over time, selection is expected to remove this sexually concordant variation; in contrast, variants with sexually antagonistic fitness effects are expected to persist longer and make up an increasingly larger proportion of the genetic variation, resulting in a less positive, or more negative, *r*_*mf*_ as the population adapts (Long et al. 2012; Connallon and Clark 2013). We do not know where our population is in this process.

An additional issue to consider is that, following the 50 generations of recombination (and lab adaptation), the RILs of the DSPR were created by 25 generations of fullsib inbreeding. During this process, there could have been some purging of variants with recessive deleterious effects. Variants with deleterious effects in both sexes would tend to be purged, reducing the amount of sexually concordant variation. Among sexually antagonistic variants, those with large recessive deleterious effects in females may be preferentially purged because the establishment of RILs depends more strongly on female than male fitness (Grieshop et al. 2017). However, without a series of additional assumptions, it is difficult to predict the net effect of the inbreeding process on fitness variation from sexually concordant versus antagonistic variants and, thereby, the effect on *r*_*mf*_.

### Quantitative Genetics Analysis

Our quantitative genetics analysis recovers significant levels of additive genetic variance for fitness in both sexes in the two mating environments where we assayed adult fitness. The heritabilities and evolvabilities of fitness were modest in both sexes in the two mating environments. These results are similar in magnitude as reported in other laboratory populations of *Drosophila* (e.g., Fowler et al. 1997; Gardner et al. 2005; Pischedda and Chippindale 2006; Long et al. 2009; Ruzicka et al. 2019) and other insect species (e.g., Archer et al. 2013; Berger et al. 2014) but our estimates of the heritability are much higher than those reported from studies of wild populations (e.g., Gustafsson 1986; Kruuk et al. 2000; Merilä and Sheldon 2000; McCleery et al. 2004; Teplitsky et al. 2009; Mcfarlane et al. 2014). Low heritability estimates in wild populations are likely the result of high levels environmental variance in fitness (Price and Schluter 1991; Houle 1992) rather than a lack of genetic variation. As noted above, the population used here is not expected to be exquisitely adapted to these assay conditions. Thus, there may be more fitness-affecting variants segregating at intermediate frequency relative to an adapted population where directional selection is expected to have reduced 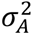.

The intersexual genetic correlation for fitness *r*_*mf*_ gives us a genome-wide summary of the extent to which the shared genome constrains the sexes from reaching their sex-specific trait optima (Lande 1980) at least in the short term. Our estimates of *r*_*mf*_ were positive in both mating environments (Figure 1), implying that, on average, genetic variants have sexually concordant fitness effects and, as a result, evolution would not be strongly constrained from improving the fitness of both sexes simultaneously in either of these environments. Though estimates for *r*_*mf*_ in both environments are significantly greater than 0, they are also far from 1. This intermediate value for *r*_*mf*_ suggests that while sexually concordant variants may be predominant, other types of variants (e.g., sex-limited and/or sexually antagonistic variants) are likely also segregating. This view is consistent with our genomic association mapping analysis: the Manhattan plots (Figure 3) suggest that sexually antagonistic genomic regions are present and are only modestly less frequent than sexually concordant ones (though this inference should be regarded with caution because of limited power). It is plausible that our estimates of positive *r*_*mf*_ reflect an overall concordance of selection as a consequence of this population being relatively maladapted to the laboratory conditions in which we measured fitness, which was not identical to how the lab population was originally created by King et al. (2012a,b).

Our significantly positive estimates of *r*_*mf*_ stands in contrast to a classic study using the same species by Chippindale et al. (2001) who reported a significantly negative *r*_*mf*_. That study measured fitness in a lab context to which their fly population was “well adapted.” However, not all studies neatly fit within the narrative in which the degree of adaptedness explains variation in *r*_*mf*_. Sharp and Agrawal (2018) estimated a positive though non-significant values for *r*_*mf*_ using another *D. melanogaster* population that was “well-adapted” to the lab in which assay conditions were similar to but did not exactly replicate maintenance conditions. A particularly interesting case is that of Collet et al. (2016) who measured *r*_*mf*_ in two descendant populations of the fly population studied by Chippindale et al. (2001). Despite being maintained and assayed similarly, the two descendant populations had significantly different values of *r*_*mf*_, with *r*_*mf*_ significantly negative in one and non-significantly positive in the other. That study demonstrates that *r*_*mf*_ can evolve substantially over a few hundred generations in ways that are not captured in any obvious way by “adaptedness”.

In addition to evolutionary changes in *r*_*mf*_ (i.e., changes in the frequencies of sequence variants), *r*_*mf*_ can be directly affected by the environment. Studies have found that *r*_*mf*_ may be highly sensitive to the ecological context in which genetic variation is expressed (e.g., Berger et al. 2014; Punzalan et al. 2014; Duffy et al. 2019). However, our estimates of *r*_*mf*_ in the two mating environments are reasonably similar (and confidence intervals are broadly overlapping). Given that sexual interactions are quite different in these two types of environments (Yun et al. 2017), it was not obvious from the outset that the two *r*_*mf*_ estimates would be so similar.

The similarity of *r*_*mf*_ estimates would be unsurprising if the genetic architecture of fitness for each sex was the same across the two environments (i.e., if *r*_*env*_ was close to 1 for both sexes). Though *r*_*env*_ was significantly positive is was also substantially less than 1 in both sexes, indicating that the genetic architecture for fitness differs somewhat across environments for both sexes. This is not surprising given that these two environments likely select for different phenotypic traits in both sexes. We have strong reasons to believe this for males. In a different experiment, populations that evolved for a few tens of generations in environments similar to those examined here had diverged significantly for male fitness in a genotype-x-environment dependent manner (Yun et al. 2019), presumably because these alternative mating environments modulate the opportunity for male harassment to result in successful mating (Yun et al. 2017). In contrast to males, females showed much weaker signs of divergence following adaptation to these alternative environments. As such, we might have expected a higher *r*_*env*_ for females than males but instead the point estimate for males is higher (0.40 vs. 0.28).

It is notable that the intersexual genetic correlations within an environment (*r*_*mf*_) and the cross-environment genetic correlations for each sex (*r*_*env*_) are all significantly positive and of similar magnitude. As mentioned above, a simple explanation for all correlations being positive is that this population contains an excess of variants that are unconditionally deleterious in the lab environment, perhaps because of insufficient time to adapt to general lab conditions. Although particular suites of phenotypes may maximize fitness for each sex and mating environments, it is plausible that many of the same morphological, physiological and behavioural traits necessary to acquire fitness are similar between the sexes in both environments. Nonetheless, it is surprising that *r*_*mf*_ and *r*_*env*_ estimates are of similar magnitude; roughly speaking, this suggests that the differences between sexes are as large as the differences between environments with respect to segregating variation. For example, knowing that a genotype confers high fitness to males in one environment is approximately as good a predictor of conferring high fitness to females in that same environment as it is of conferring high fitness to males in the other environment.

### Association Mapping Analysis

Our association mapping analysis revealed few, if any, genomic regions with very strong statistical associations with the traits examined. Consequently, we took a more exploratory approach to evaluating whether there was any evidence of genetic associations beyond what would be expected by chance, using permutations to create appropriate null distributions. The results of these analyses were somewhat inconclusive. On the one hand, in almost all cases the observed number of associations was larger than expected. However, the permutation-based null distributions have high variance and, in only some cases, was the observed number of associations unusually high relative to the corresponding null distribution. With some criteria, there was a substantial excess of associations with SA but not SC; with other criteria, there was an excess of associations with SC but not SA.

In general, the power to detect a significant genomic association is inversely related to the minor allele frequency (MAF) at loci underlying trait variation. An advantage of this synthetic population is that the variants segregating in this panel should have high MAFs (i.e., a neutral expectation of 12.5%) compared to natural populations. However, the observed MAFs are often lower than 12.5%, presumably due to 50 generations of drift and selection. Of course, statistical power is also directly related to effect size. Because of selection, the standing variation will be depauperate of variants with large directional effects (e.g., unconditionally deleterious) segregating at high frequency. It is likely there are numerous variants that make small contributions to fitness variation, but we would not expect to detect them via this association mapping approach. Nonetheless, it seemed plausible we could detect large-effect sexually antagonistic variants if such alleles were segregating. Theory indicates that large-effect sexually antagonistic variants are more readily maintained by balancing selection than small effect ones (Kidwell et al. 1977; Connallon and Clark 2012). Our failure to observe any strong associations implies that the loci underlying fitness are unlikely to be of large effect.

Our tentative interpretation of the association study is that effect sizes are small to moderate for both SA and SC as we tend to see more modest associations than expected yet fail to find very strong associations. In addition, these data suggest that SA variation is not particularly rare relative to (detectable) SC variation as we see only modestly stronger evidence for associations with SC than SA (Table 2).

Using a permissive threshold to generate larger lists of candidate regions, we found some evidence of enriched GO terms for SA and SC in the complex but not the simple environment. In their genomic analysis of sexual antagonism, Ruzicka et al. (2019) found little evidence of enriched GO categories and those they did where not obviously linked to sexually dimorphic traits. One interpretation of these results is that sexual antagonism is diffuse affecting many types of processes and/or is especially prone to persisting in genes affecting a variety of basic functions that are under selective constraint. However, another interpretation is that studies such as ours lack power to meaningfully identify genomic regions experiencing sexually antagonistic selection. On the other hand, even though the terms identified are not readily biologically interpretable, the existence of significantly enriched GO terms for both SA and SC in the complex environment provide another form of evidence that the association analysis is not just noise.

### Mutational Burden

Examining entire classes of variants, rather than individual variants, provides greater power to detect smaller effects. We estimated that each additional LoF reduces female fitness by, on average, ∼0.7%, and each additional indel by ∼0.06%. In contrast, we find no significant associations between any of the three classes of putatively deleterious variants and relative fitness in males in either mating environment. The results imply that selection acting on mutational burden is stronger in females than in males.

These results are surprising considering many studies have found evidence for stronger selection acting on variants expressed in males. Several studies have estimated the sex-specific selection acting on marker mutations in *Drosophila* (Whitlock and Bourguet 2000; Pischedda and Chippindale 2005; Stewart et al. 2005; Sharp and Agrawal 2008; Hollis et al. 2009). These studies have generally found evidence in support of the hypothesis that the strength of selection tends to be stronger in males. However, it is unclear whether results from visible marker mutations apply more broadly. Several studies examining the fitness consequences of novel mutations via mutation accumulation (MA) experiments in *Drosophila* and other species (e.g., Enders and Nunney 2010; Mallet et al. 2011; Mcguigan et al. 2011; Sharp and Agrawal 2013; Grieshop et al. 2016) have also generally supported the hypothesis of stronger selection through males. While an advantage of using MA lines is that they capture the fitness effects of more ‘typical’ mutations, one disadvantage is that the causative mutations are typically completely unknown. Fitness results from MA studies may be dominated by a small subset of the mutations that have large effects. This is problematic if such mutations are unrepresentative in other ways. For example, perhaps mutations with male-limited effects are disproportionately likely to be of large effect.

It remains unclear why our current study found evidence of stronger selection in females than males whereas past studies using phenotypic marker mutations or MA lines found the opposite. One possibility is that the genetic variation in the DSPR has been filtered by selection in the wild (i.e., which variants were present in the eight founder lines for the DSPR) in contrast to MA studies that capture the fitness effects of all novel mutations (excluding only extremely large effect variants e.g., lethal or sterilizing variants). Variants causing strong effects in the wild (in either sex) are expected to be underrepresented relative to mutational input. One possibility for the discrepancy between this study and others is that fitness effects on males are reasonably consistent between field and lab conditions whereas effects on females are more variable between environments. Thus, variants with large fitness effects in males in the lab would be underrepresented whereas those causing large female effects in the lab would not be so underrepresented due to past selection in the wild. Perhaps selection is more consistent on males between field and lab because their fitness is so heavily determined by their interactions with other flies via mate competition whereas female fitness may be more directly affected by the environment itself (e.g., temperature, humidity, food availability).

A second possibility for the discrepancy between this and past study is from differences in how fitness was assayed. First, viability is a potentially important source of fitness variation in our study but minimized in many other studies. We measured fitness of adults via offspring produced after a six day ‘mating interaction’ period whereas other fly studies typically use much shorter mating periods. Our protracted period of mating interaction allows for more fitness variation via adult mortality, especially so for females; males that die during this period can still contribute offspring via matings prior to death but females dying prior to the day we allowed for egg laying would prevent any contribution to offspring production in our study (i.e., fitness effect of mortality during the mating period is stronger on females than males).

While the type of variants we studied are often presumed to be deleterious, very few studies have attempted to validate this presumption by examining associations with fitness. Two recent studies have employed similar methods to investigate the aggregate fitness consequences of putatively deleterious variants. Yang et al (2017) employed a comparatively conservative criteria to filter for putatively deleterious variants in maize (*Zea mays*) by filtering SNPs based on both their genomic evolutionary rate profiling (GERP) scores—a measure of evolutionary constraint on a SNP (Cooper et al. 2005)—and the predicted functional consequence of variants. Brown and Kelly (2019) took a more liberal approach to identify deleterious variants in yellow monkeyflower (*Mimulus guttatus*) by filtering out any variant with a minor allele cut-off greater than 5%. Both studies we able to identify significant associations between mutation burden and measures of fitness. Our own study did not provide evidence of negative selection when we averaged over both sexes. For LoFs and indels, we detected negative selection in females but not males. For radical amino acid missense variants, sex-average selection was positive. The latter class of variant is the one for which we have the weakest *a priori* expectation for deleterious effects. Our results suggest that some of these variants may be beneficial at least in the lab environment.

### Conclusion

Any study of the genetic variation in fitness should be considered with respect to the history of the population in relation to the environment in which fitness was assayed. Our study likely represents a modestly, but not exquisitely, adapted population. Relative to an adapted population at equilibrium in a constant environment, we would expect the genetic lines used here to harbour more sexually concordant variation. Despite this expected “bias” towards concordance, both the quantitative genetic and association mapping results suggest sex differences in the genetic architecture of fitness. The observed intersexual genetic correlations are greater than zero, yet they are also substantially less than 1. Within the limits of our statistical power, there is evidence for genomic regions that are sexually concordant but also regions that are sexually antagonistic. While the history of the population is expected to affect the relative amounts of concordant versus antagonistic variation, there is no *a priori* reason it should differentially affect the sexes with respect to the fitness effects of putatively deleterious variants. Nonetheless, we find evidence of stronger selection in females than males, contrary to studies of selection against phenotypic markers and new mutations. While caution should be used in interpreting these results, they add to the existing empirical evidence of sex differences in selective effects.

## Acknowledgements

We are grateful to many undergraduate assistants who helped with data collection, especially Nina Adler, Malak Bayoumi, Elenore Breslow, and Matthew Lindsay. We thank Karl Grieshop, Julia Kreiner, George Sandler, and Santiago Sánchez-Ramírez for statistics/bioinformatics help and advice. DSPR lines were generously provided by Stuart MacDonald. Funding for this research was provided in the form of an Ontario Graduate Scholarship (OGS) to AS and an NSERC Discovery Grant to AFA.

## Notes

### Competing Interest Statement

The authors have declared no competing interest.

### Summary of Updates

Changes to analyses and presentation. Major results remain the same.

