## Supplementary Material for "An investigation of the sex-specific genetic architecture of fitness in *Drosophila melanogaster*"

Table of Contents


**Supplemental Text:**

1. Quantitative genetics of fitness measured from a group of individuals
2. Quantitative genetics of SC and SA.
3. Determining mutational burden of each DSPR RIL.

**Supplemental Figures:**

1. Schematic of fitness assays.
2. Graphical representation of SA and SC axes.
3. Radical amino acid burden of 357 DSPR RILs.
4. Manhattan plots for male and female fitness.

**Supplemental Tables:**

1. Counts of DSPR RILs with sex-specific fitness estimates in either, both or neither simple or complex mating environments.
2. The number haplotype blocks associated with SA and/or SC passing a specified *q*-value criterion.
3. The number haplotype blocks associated with female and/or male fitness passing a specified statistical criterion.
4. The number haplotype blocks associated with female and/or male fitness passing a specified *q*-value criterion.
5. Gene Ontology (GO) enrichment for SA in the simple mating environment.
6. Gene Ontology (GO) enrichment for SC in the simple mating environment.
7. Gene Ontology (GO) enrichment for SA in the complex mating environment.
8. Gene Ontology (GO) enrichment for SC in the complex mating environment.
9. Gene Ontology (GO) enrichment for female relative fitness in the simple mating environment.
10. Gene Ontology (GO) enrichment for male relative fitness in the simple mating environment.
11. Gene Ontology (GO) enrichment for female relative fitness in the complex mating environment.
12. Gene Ontology (GO) enrichment for male relative fitness in the complex mating environment.

**a.**

**Supplemental Text 1**: *Quantitative genetics of fitness measured from a group of individuals*

We estimated the fitness of groups consisting of four focal individuals, all of whom share a haploid genome from focal RIL $r$. Each individual has its own unique other haploid genome that has come at random from an outbred population. The genotypic value of the $i^{\text{th}}$ individual in the $j^{\text{th}}$ replicate from focal RIL $r$ is

$$G_{ijr} = A_{ijr} + D_{ijr}$$

where $A$ and $D$ represent the additive and dominance values respectively. The additive genetic value can be decomposed as

$$A_{ijr} = a_{fr} + a_{o,ijr}$$

where $a_{fr}$ is the sum of the average effect of alleles received from the focal RIL parent (i.e., RIL $r$) and $a_{o,ijr}$ is the sum of the average effect of alleles from the other parent. Note that all individuals derived from RIL $r$ share the same value of $a_{fr}$ whereas each individual's value of $a_{o,ijr}$ can be unique. Moreover, the $a_o$ values are uncorrelated among individuals with the same RIL parent and to the $a_r$ values.

The phenotypic value of the $i^{\text{th}}$ individual in the $j^{\text{th}}$ replicate from focal RIL $r$ is

$$Z_{ijr} = G_{ijr} + E_{ijr}$$

where $E_{ijr}$ is the environmental value that can be further decomposed to $E_{ijr} = E_{s,ijr} + E_{c,ijr}$ where $E_s$ represents environmental effects specific to a single individual and $E_c$ represents the common environmental effect experienced by each of the focal individuals in the same replicate.

The total success of the group of focal individuals within a replicate (i.e., replicate $j$ from RIL $r$) is assumed to be the sum of the four individual phenotypic values:

$$Y_{jr} = \sum_{i=1}^{4} Z_{ijr}$$

$$= 4a_{f,r} + 4E_{c,jr} + \sum_{i=1}^{4} a_{o,ijr} + \sum_{i=1}^{4} D_{ijr} + \sum_{i=1}^{4} E_{s,ijr}$$

Biologically, the implicit assumption in doing the sum above is that focal individuals do not interfere with one another. While this assumption is imperfect, each replicate in our study contained only four focal individuals and 12 competitors, so that most fitness gain of a focal individual should come at the expense of a competitor.

The total variance among groups (i.e., the unit of observation; *Y*) is

$$\sigma_Y^2 = 16\sigma_{af}^2 + 16\sigma_{Ec}^2 + 4\sigma_{ao}^2 + 4\sigma_D^2 + 4\sigma_{Es}^2$$

Noting that $\sigma_{af}^2 = \sigma_{ao}^2 = \frac{\sigma_A^2}{2}$, where $\sigma_A^2$ is the additive genetic variance in fitness, then,

$$\sigma_Y^2 = 10\sigma_A^2 + 16\sigma_{Ec}^2 + 4\sigma_D^2 + 4\sigma_{Es}^2$$

It seems probable that $\sigma_{Ec}^2 \ll \sigma_{Es}^2$ because fitness of the group is measured as a proportion of offspring when focal individuals compete against a reasonably large number of competitors. In this case,

$$\sigma_Y^2 = 10\sigma_A^2 + 4\sigma_D^2 + 4\sigma_{Es}^2$$

Environmental effects are not shared among replicates of the same RIL, nor are $a_o$ values or dominance deviations. Only the additive genetic effects from the RIL itself (i.e., $a_{fr}$ values) cause replicates of the same RIL to covary. Thus, the variance among RILs is $\sigma_{RIL}^2 = 16\sigma_{af}^2 = 8\sigma_A^2$. The variance unexplained by RIL—the residual variance—is $\sigma_e^2 = \sigma_Y^2 - \sigma_{RIL}^2 = 2\sigma_A^2 + 4\sigma_D^2 + 4\sigma_{Es}^2$ Using this, we can estimate heritability as

$$h^2 = \frac{\sigma_{RIL}^2/2}{\sigma_Y^2 - \left(\frac{3}{4}\right)\sigma_{RIL}^2} = \frac{\sigma_{RIL}^2}{2\left(\sigma_e^2 + \frac{\sigma_{RIL}^2}{4}\right)}$$

as this is approximately

$$h^2 \approx \frac{\sigma_A^2}{\sigma_A^2 + \sigma_D^2 + \sigma_{Es}^2}$$

We can estimate evolvability as

$$e_\mu = \sigma_{RIL:j}^2/\mu_j^2$$

**Supplemental Text 2**: *Quantitative genetics of SC* and *SA*

For a *RIL* with sex-specific relative fitness values $w_f$ and $w_m$, its corresponding values for *SC* and *SA* are given by

$$SA = (w_f - w_m)/\sqrt{2} \quad \text{(S2.1a)}$$

and

$$SC = (w_f + w_m)/\sqrt{2} \quad \text{(S2.1b)}$$

Because *SA* and *SC* are linear transformations of $w_f$ and $w_m$, the variances of *SA* and *SC* are simple functions of the (co)variances of $w_f$ and $w_m$:

$$V(SA) = \frac{V(w_f)}{2} + \frac{V(w_m)}{2} - C(w_f, w_m) \quad \text{(S2.2a)}$$

and

$$V(SC) = \frac{V(w_f)}{2} + \frac{V(w_m)}{2} + C(w_f, w_m) \quad \text{(S2.1b)}$$

The values above represent phenotypic variances for all 4 traits ($w_f$, $w_m$, *SA*, *SC*) but the same logic would lead to equivalent expressions for the additive genetic variances of *SA* and *SC* as functions of the additive genetic variances $w_f$ and $w_m$ and their additive genetic covariance.

One could estimate a measure equivalent to "heritability" as

$$h^2(SA) = \frac{\left(\frac{G_{ff}}{2} + \frac{G_{mm}}{2} - G_{fm}\right)}{\frac{V(w_f)}{2} + \frac{V(w_m)}{2} - C(w_f, w_m)} \quad \text{(S2.3a)}$$

$$h^2(SC) = \frac{\left(\frac{G_{ff}}{2}+\frac{G_{mm}}{2}+G_{fm}\right)}{\frac{V(w_f)}{2}+\frac{V(w_m)}{2}+C(w_f,w_m)} \tag{S2.3b}$$

where *G* terms represent additive genetic (co)variances for relative fitness.

Note that standard traits (e.g., body size or fitness) are measured at the individual level and an individual's trait value is composed of that individual's genetic and environmental values. In contrast, *SA* and *SC* traits do not exist at the individual level. Environmental effects contribute to variance in *SA* and *SC* through $V(w_f)$ and $V(w_m)$— the first two terms in equation S2.2—but not through $C(w_f, w_m)$ because fitness is measured independently in the two sexes. Thus, for calculating heritabilities of *SA* and *SC*, we assumed $C(w_f, w_m) = G_{fm}$.

**Supplemental Text 3:** *Determining mutational burden of each DSPR RIL*

Genomic Imputation of Inbred Genotypes

To impute SNPs segregating among inbred genotypes, we used the results of a HMM implemented to uncover the parental ancestry of each genotype at regularly spaced 10kb intervals along the genome of each inbred genotype and a table of SNPs segregating in each parental line from the DSPR webpage (http://wfitch.bio.uci.edu/~dspr/). We filtered out all sites in the parental SNP table where less than 95% of SNP calls were to either the reference or alternate allele in each of the parental lines, sites with less than 15 reads and all multiallelic sites. Within each line, we filtered out any 10kb intervals where the ancestry assignment probability of the most probable of the eight parental lines was less than 95%. This filtering step removed an average of ~8.5% of blocks from each RIL.

Imputation of insertion/deletion (indel) variants began by obtaining indel VCFs for each of the eight DSPR parental lines from the *Drosophila* Genome Nexus (DGN) webpage (https://www.johnpool.net/genomes.html). We filtered out indels not supported by at least 30 reads. We then imputed indels in the 357 DSPR lines in the same manner as described above for SNPs.

*Identifying putatively rare variants from the DSPR*

Because deleterious variants are expected to segregate at low frequency at mutation-selection balance, a reasonable approach to identifying putatively deleterious variants might include focusing only on low frequency variants. To this end, we filtered out any variants segregating in the DSPR that were also segregating in two geographically disparate outbred populations of *D. melanogaster*: 197 genotypes from Zambia (*Drosophila* Population Genomic

Project Phase 3 [DPGP3]) and 205 genotypes from Raleigh, North Carolina (*Drosophila* Genetic Reference Panel [DGRP]).

FASTA files containing whole genome sequences for all individuals sampled in both the DPGP3 and DGRP populations were obtained from the DGN webpage. These genome sequences were generated following a standardized alignment and quality control procedures described elsewhere (Lack et al. 2015, 2016). Using Perl scripts supplied on the DGN webpage, we further filtered sites that showed evidence of high relatedness and recent admixture. We concatenated all genomes in a single population into a single multi-FASTA alignment file and generated a VCF containing SNPs for each population using *SNP-sites* (Page et al. 2016). Next, we filtered out any SNPs in the DSPR that were present in VCFs for either of the two outbred populations (i.e., sites segregating for variants in the DSPR that were also captured as segregating in the two outbred population samples were excluded from downstream analyses).

Next, we needed to determine which variant in the DSPR was the major allele in nature. We chose one genome at random from the DPGP3 (ZI-103) and one genome at random from the DGRP (RAL-100). For both genomes, we filtered out any sites that were polymorphic within the populations that they were sampled from and then filtered out any sites that differed between these two sequences including all sites that were masked in either sequence (these sites were also filtered out from the DSPR). We compared this sequence (which we term the 'outbred consensus sequence') back to the DSPR VCF file described above and filtered out any sites where one of the two segregating alleles in the DSPR was not identical to the outbred consensus sequence. Finally, for each site in the DSPR, we deemed the variant matching the outbred consensus sequence as the major allele.

Indels were filtered from DSPR lines in a similar manner. We obtained indel VCFs for both outbred populations from the DGN webpage and combined each individual VCF using the '*vcf-merge*' command in *VCFtools* (Danecek et al. 2011). Finally, we filtered out any indels in the DSPR lines that shared a starting coordinate with indels segregating in the either the DGRP or DPGP3 indel VCFs.

*Functional annotation of genomic variants in the DSPR and identifying putatively rare deleterious variants*

We predicted the functional consequences of minor allele SNPs and indels in the DSPR lines using VEP (McLaren et al. 2010). Genomic coordinates for all VCFs were updated to release 6 of the Drosophila reference genome using the UCSC liftOver tool (Kent et al. 2002) prior to passing VCFs to VEP. We retained only variants that fell within coding regions. For SNPs, we focused on two classes of variants: those variants labelled by VEP as high 'IMPACT' and missense mutations resulting in radical amino acid changes. Variants with high IMPACT scores include variants predicted to have severe functional consequences including splice acceptor/donor disrupting variants and stop loss/gain and start loss variants. Following Sohail et al. (2017), we refer to this combined set of high IMPACT variants as 'loss-of-function' (LoF) mutations. For each missense variant we characterized the resulting amino acid change as either a conservative or radical change based on an amino acid classification scheme proposed by Zhang (2000) that classifies amino acids into six categories that vary with respect to polarity and volume. An amino acid change is deemed 'radical' if the major and minor alleles code for amino acids that are in different categories with respect to either polarity or volume. Radical substitutions are under stronger purifying selection in *Drosophila* than conservative substitutions (Smith 2003). Finally, we use VEP to map indels to the reference genome and filtered out all indels that did not map to coding regions.

### Supplemental Figures

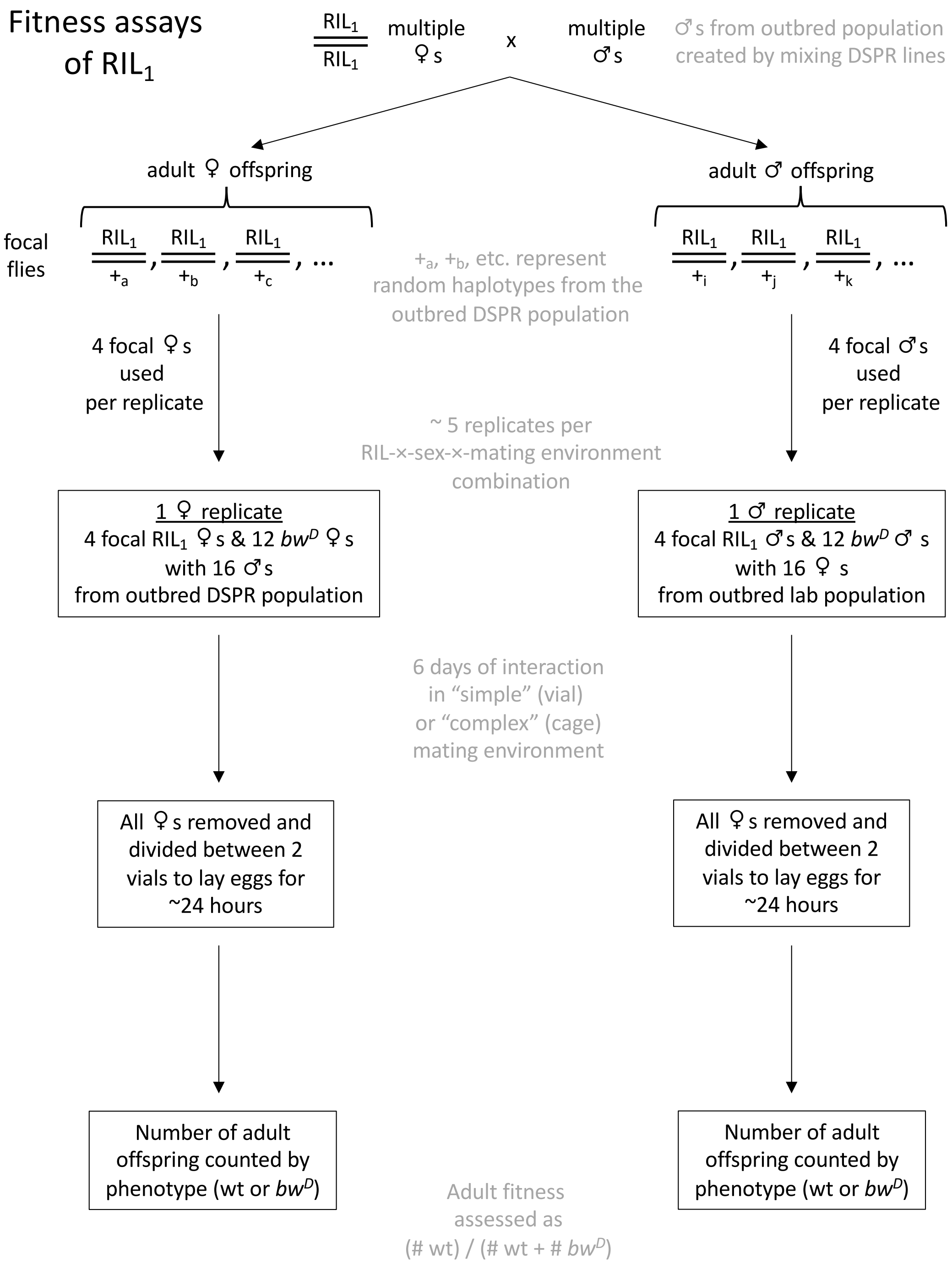


**Figure S1.** A schematic of the fitness assays. The procedure is outlined with respect to one focal haplotype, "$RIL_1$". In the top two steps, diploid genotypes are depicted as a combination of two haplotypes. (X and Y chromosomes are not shown but note focal flies of both sexes receive an X from $RIL_1$). See main text for details.

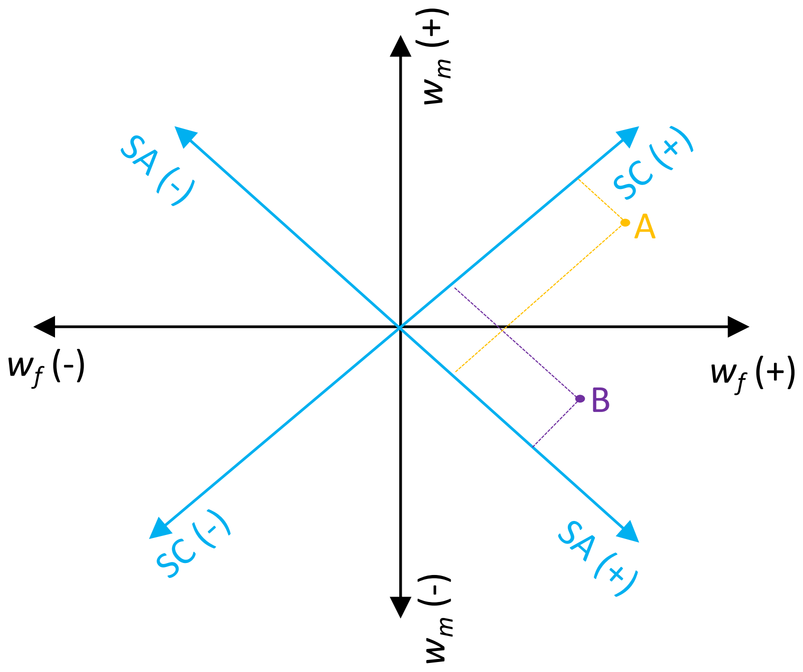


**Figure S2.** Graphical representation of SA and SC axes. The SC and SA axes (shown in blue) are a 45° rotation from the female ($w_f$) and male ($w_m$) relative fitness axes. The origin with respect to $w_f$ and $w_m$ is intended to represent a genotype with average relative fitness with respect for both sexes. Two points are shown as examples. Point A represents a haplotype that has higher than average fitness with respect to both sexes, but more so for female fitness. It's values with respect to SC and SA are depicted by the dashed lines orthogonal to the SC and SA axes; it has a strongly positive SC value and weakly positive SA value. Point B represents a haplotype that has higher than average female fitness but lower than average male fitness. This corresponds to weakly positive SC value and a more strongly positive SA value.

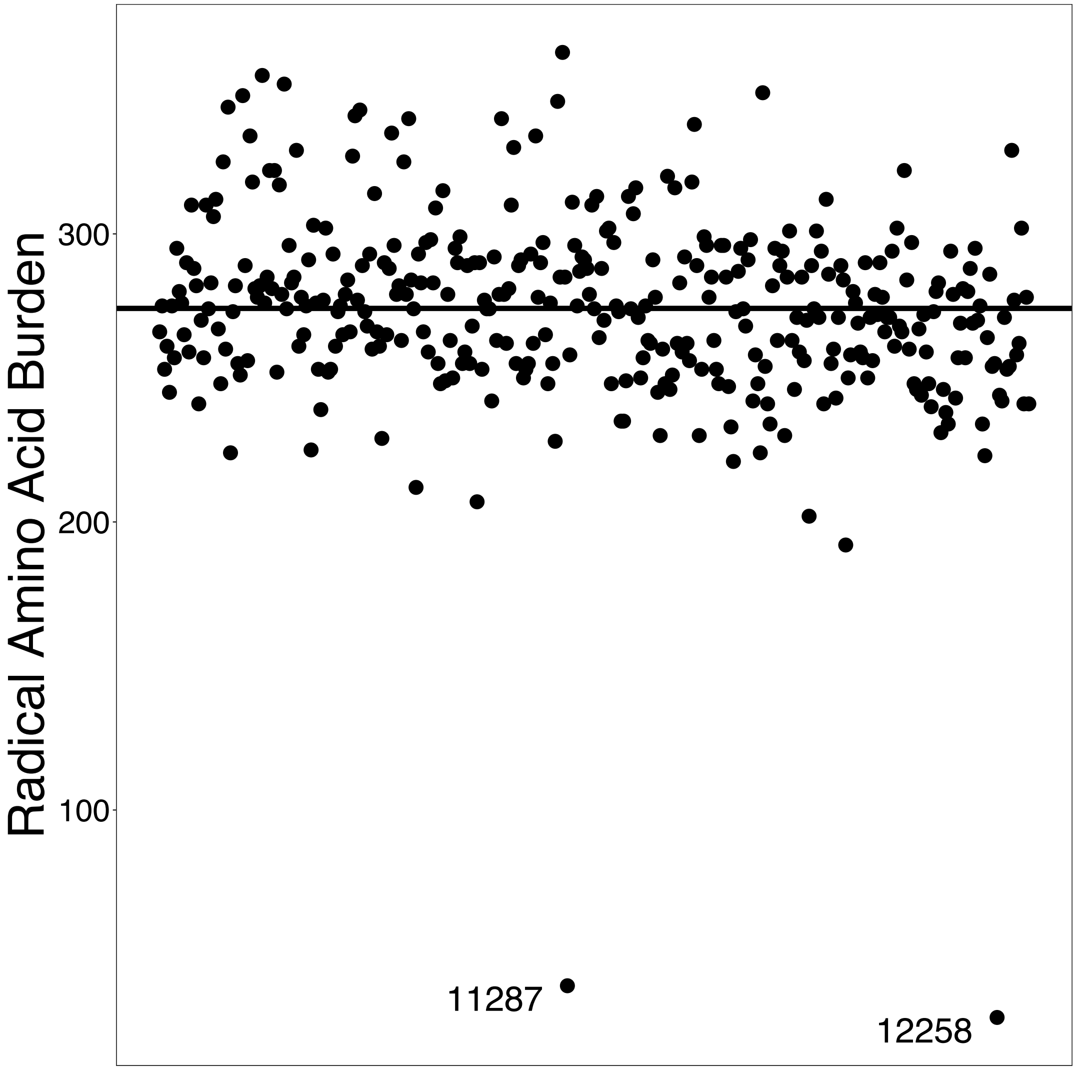


**Figure S3:** Radical amino acid burden of 357 DSPR RILs. Solid line indicates mean radical amino acid burden of all RILs in the sample. Labelled points are RILs that were deemed to be outliers based on visual inspection and were removed from further analysis.

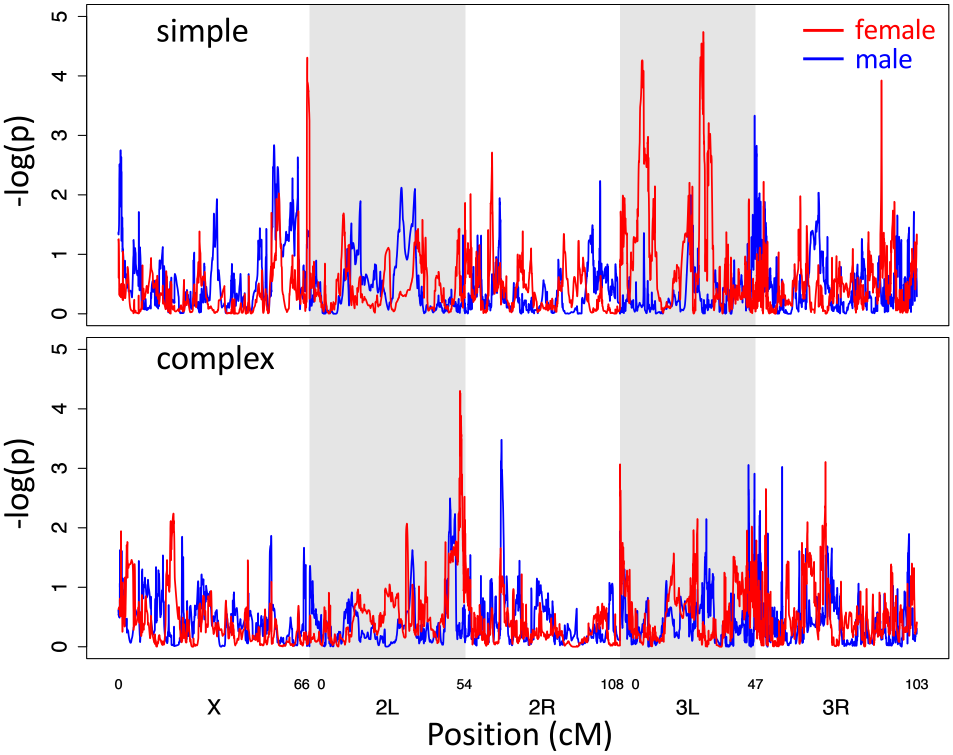


**Figure S4.** Manhattan plots for male and female fitness. Strength of statistical support for associations between 10kb haplotype blocks along the genome with female (red) and male (blue) relative fitness in DSPR lines when measured in the simple (top) or complex (bottom) mating environment.

**Supplemental Tables**

**Table S1:** Counts of DSPR RILs with sex-specific fitness estimates in either, both or neither simple or complex mating environments.

| | | **Male Fitness** | | | |
|---|---|---|---|---|---|
| | | **Simple and Complex Mating Environments** | **Simple Mating Environment Only** | **Complex Mating Environment Only** | **No Estimates** |
| **Female Fitness** | **Simple and Complex Mating Environments** | 300 | 5 | 3 | 0 |
| | **Simple Mating Environment Only** | 3 | 34 | 0 | 2 |
| | **Complex Mating Environment Only** | 3 | 0 | 7 | 0 |
| | **No Estimates** | 0 | 0 | 0 | 0 |

**Table S2:** The number haplotype blocks associated with SA and/or SC passing a specified $q$-value criterion. For this analysis, only every 7th haplotype block was tested (1682 blocks tested). $n$ is the observed number in the real (i.e., unpermuted data). $\bar{n}_{perm}$ is the mean number of associations in permutated data sets. $P_{perm}$ is the proportion of permutations with as many as or more than the observed number of associations ($n$); cases where the observed value is the upper 5% of the distribution (i.e., $P_{perm} < 0.05$) are shown in bold.

| Criterion for 'significant' association | Metric | Any trait | SA | SC | SA in simple environment | SC in simple environment | SA in complex environment | SC in complex environment |
|---|---|---|---|---|---|---|---|---|
| | $n$ | 0 | 0 | 15 | 0 | 5 | 0 | 12 |
| $q \leq 0.1$ | $\bar{n}_{perm}$ | 1.0 | 1.3 | 1.5 | 2.3 | 2.0 | 2.0 | 2.6 |
| | $P_{perm}$ | 1 | 1 | **0.03** | 1 | 0.058 | 1 | 0.047 |
| | $n$ | 25 | 0 | 40 | 252 | 109 | 0 | 17 |
| $q \leq 0.3$ | $\bar{n}_{perm}$ | 10.3 | 16.1 | 17.0 | 35.4 | 45.1 | 38.8 | 34.8 |
| | $P_{perm}$ | 0.095 | 1 | 0.111 | **0.037** | 0.089 | 1 | 0.169 |

**Table S3:** The number haplotype blocks associated with female and/or male fitness passing a specified statistical criterion. For this analysis, only every 7th haplotype block was tested (1682 blocks tested). $n$ is the observed number in the real (i.e., unpermuted data). $\bar{n}_{perm}$ is the mean number of associations in permutated data sets. $P_{perm}$ is the proportion of permutations with as many as or more than the observed number of associations ($n$); cases where the observed value is the upper 5% of the distribution (i.e., $P_{perm} < 0.05$) are shown in bold.

| Criterion for 'significant' association | Metric | Any trait | Female fitness | Male fitness | Female fitness in simple environment | Male fitness in simple environment | Female fitness in complex environment | Male fitness in complex environment |
|---|---|---|---|---|---|---|---|---|
| | $n$ | 22 | 14 | 8 | 14 | 1 | 0 | 7 |
| $p_{crit}$ = 0.001 | $\bar{n}_{perm}$ | 7.8 | 4.2 | 3.6 | 2.4 | 1.8 | 1.8 | 1.8 |
| | $P_{perm}$ | 0.076 | 0.075 | 0.155 | **0.039** | 0.355 | 1 | 0.086 |
| | $n$ | 154 | 78 | 76 | 36 | 62 | 42 | 14 |
| $p_{crit}$ = 0.01 | $\bar{n}_{perm}$ | 72.0 | 36.5 | 35.6 | 19.0 | 17.5 | 17.5 | 18.1 |
| | $P_{perm}$ | **0.031** | 0.073 | 0.065 | 0.141 | 0.026 | 0.086 | 0.484 |
| | $n$ | 792 | 285 | 507 | 113 | 318 | 172 | 189 |
| $p_{crit}$ = 0.05 | $\bar{n}_{perm}$ | 354 | 177 | 177 | 90.1 | 87.3 | 86.7 | 89.5 |
| | $P_{perm}$ | **<0.001** | 0.063 | **<0.001** | 0.256 | **0.001** | 0.052 | **0.036** |
| | $n$ | 1174 | 505 | 669 | 220 | 386 | 285 | 283 |
| $p_{crit}$ = 0.1 | $\bar{n}_{perm}$ | 700 | 350 | 350 | 176 | 173 | 174 | 177 |
| | $P_{perm}$ | **0.001** | 0.066 | **<0.001** | 0.244 | **0.004** | 0.072 | 0.068 |

**Table S4:** The number haplotype blocks associated with female and/or male fitness passing a specified $q$-value criterion. For this analysis, only every 7th haplotype block was tested (1682 blocks tested). $n$ is the observed number in the real (i.e., unpermuted data). $\bar{n}_{perm}$ is the mean number of associations in permutated data sets. $P_{perm}$ is the proportion of permutations with as many as or more than the observed number of associations ($n$); cases where the observed value is the upper 5% of the distribution (i.e., $P_{perm} < 0.05$) are shown in bold.

| Criterion for 'significant' association | Metric | Any trait | Female fitness | Male fitness | Female fitness in simple environment | Male fitness in simple environment | Female fitness in complex environment | Male fitness in complex environment |
|---|---|---|---|---|---|---|---|---|
| | $n$ | 10 | 14 | 0 | 14 | 0 | 0 | 6 |
| $q \leq 0.1$ | $\bar{n}_{perm}$ | 1.6 | 2.2 | 1.1 | 2.9 | 1.9 | 2.1 | 3.1 |
| | $P_{perm}$ | **0.036** | **0.039** | 1 | 0.052 | 1 | 1 | 0.063 |
| | $n$ | 43 | 42 | 507 | 47 | 333 | 24 | 7 |
| $q \leq 0.3$ | $\bar{n}_{perm}$ | 10.5 | 20.0 | 13.9 | 49.0 | 43.3 | 34.7 | 38.5 |
| | $P_{perm}$ | 0.059 | 0.093 | **0.004** | 0.128 | **0.035** | 0.135 | 0.213 |

**Table S5:** Gene Ontology (GO) enrichment for SA in the simple mating environment. GO categories enriched for genes annotated within 5kb of the leading haplotype block in each statistically significant genomic cluster (see Methods).

| Gene Ontology (GO) | GO Term | Description | *p*-value | *q*-value |
|---|---|---|---|---|
| **Biological Process** | GO:0034220 | Ion transmembrane transport | < 0.001 | 1.00 |
| | GO:0055085 | Transmembrane transport | < 0.001 | 1.00 |
| | GO:0032501 | Multicellular organismal process | < 0.001 | 1.00 |
| | GO:0098916 | Anterograde trans-synaptic signaling | < 0.001 | 1.00 |
| | GO:0007268 | Chemical synaptic transmission | < 0.001 | 1.00 |
| | GO:0099537 | Trans-synaptic signaling | < 0.001 | 1.00 |
| **Molecular Function** | GO:0015318 | Inorganic molecular entity transmembrane transporter activity | < 0.001 | 1.00 |
| | GO:0015075 | Ion transmembrane transporter activity | < 0.001 | 1.00 |
| | GO:0022848 | Acetylcholine-gated cation-selective channel activity | < 0.001 | 0.794 |
| **Cellular Component** | GO:0032809 | Neuronal cell body membrane | < 0.001 | 0.191 |
| | GO:0044298 | Cell body membrane | < 0.001 | 0.095 |
| | GO:0031224 | Intrinsic component of membrane | < 0.001 | 0.227 |
| | GO:0045202 | Synapse | < 0.001 | 0.263 |
| | GO:0005892 | Acetylcholine-gated channel complex | < 0.001 | 0.225 |

**Table S6:** Gene Ontology (GO) enrichment for SC in the simple mating environment. GO categories enriched for genes annotated within 5kb of the leading haplotype block in each statistically significant genomic cluster (see Methods).

| Gene Ontology (GO) | GO Term | Description | *p*-value | *q*-value |
|---|---|---|---|---|
| **Biological Process** | GO:0090251 | Protein localization involved in establishment of planar polarity | < 0.001 | 1 |
| **Molecular Function** | GO:0008239 | Dipeptidyl-peptidase activity | < 0.001 | 0.105 |
| **Cellular Component** | - | - | - | - |

**Table S7**: Gene Ontology (GO) enrichment for SA in the complex mating environment. GO categories enriched for genes annotated within 5kb of the leading haplotype block in each statistically significant genomic cluster (see Methods). Bold font indicates GO term is significant at FDR *q*-value < 0.05.

| Gene Ontology (GO) | GO Term | Description | *p*-value | *q*-value |
|---|---|---|---|---|
| Biological Process | **GO:0004656** | **Procollagen-proline 4-dioxygenase activity** | **< 0.001** | **0.004** |
| | **GO:0004656** | **Peptidyl-proline 4-dioxygenase activity** | **< 0.001** | **0.002** |
| | **GO:0031543** | **Peptidyl-proline dioxygenase activity** | **< 0.001** | **0.001** |
| | **GO:0019798** | **Procollagen-proline dioxygenase activity** | **< 0.001** | **0.001** |
| | **GO:0016702** | **Oxidoreductase activity, acting on single donors with incorporation of molecular oxygen, incorporation of two atoms of oxygen** | **< 0.001** | **0.002** |
| | **GO:0016701** | **Oxidoreductase activity, acting on single donors with incorporation of molecular oxygen** | **< 0.001** | **0.003** |
| | **GO:0016706** | **Oxidoreductase activity, acting on paired donors, with incorporation or reduction of molecular oxygen, 2-oxoglutarate as one donor, and incorporation of one atom each of oxygen into both donors** | **< 0.001** | **0.006** |
| | **GO:0051213** | **Dioxygenase activity** | **< 0.001** | **0.012** |
| | **GO:0031418** | **L-ascorbic acid binding** | **< 0.001** | **0.021** |
| | GO:0043177 | Organic acid binding | < 0.001 | 0.107 |
| | GO:0031406 | Carboxylic acid binding | < 0.001 | 0.097 |
| | GO:0048029 | Monosaccharide binding | < 0.001 | 0.142 |
| Molecular Function | GO:0015318 | Inorganic molecular entity transmembrane transporter activity | < 0.001 | 1.00 |
| | GO:0015075 | Ion transmembrane transporter activity | < 0.001 | 1.00 |
| | GO:0022848 | Acetylcholine-gated cation-selective channel activity | < 0.001 | 0.794 |
| Cellular Component | - | - | - | - |

**Table S8:** Gene Ontology (GO) enrichment for SC in the complex mating environment. GO categories enriched for genes annotated within 5kb of the leading haplotype block in each statistically significant genomic cluster (see Methods). Bold font indicates GO term is significant at FDR *q*-value < 0.05.

| Gene Ontology (GO) | GO Term | Description | *p*-value | *q*-value |
|---|---|---|---|---|
| **Biological Process** | GO:0019731 | Antibacterial humoral response | < 0.001 | 0.117 |
| | GO:0019730 | Antimicrobial humoral response | < 0.001 | 0.4676 |
| | GO:0006959 | Humoral immune response | < 0.001 | 0.638 |
| | GO:0050830 | Defense response to Gram-positive bacterium | < 0.001 | 0.653 |
| **Molecular Function** | **GO:0008239** | **Dipeptidyl-peptidase activity** | **< 0.001** | **0.008** |
| | GO:0004177 | Aminopeptidase activity | < 0.001 | 0.444 |
| | GO:0003690 | Double-stranded DNA binding | < 0.001 | 0.718 |
| | GO:0003677 | DNA binding | < 0.001 | 0.645 |
| **Cellular Component** | GO:0005634 | Nucleus | < 0.001 | 0.809 |

**Table S9:** Gene Ontology (GO) enrichment for female relative fitness in the simple mating environment. GO categories enriched for genes annotated within 5kb of the leading haplotype block in each statistically significant genomic cluster (see Methods).

| Gene Ontology (GO) | GO Term | Description | *p*-value | *q*-value |
|---|---|---|---|---|
| **Biological Process** | - | - | - | - |
| **Molecular Function** | GO:0004180 | Carboxypeptidase activity | < 0.001 | 1.00 |
| **Cellular Component** | - | - | - | - |

**Table S10:** Gene Ontology (GO) enrichment for male relative fitness in the simple mating environment. GO categories enriched for genes annotated within 5kb of the leading haplotype block in each statistically significant genomic cluster (see Methods). Bold font indicates GO term is significant at FDR *q*-value < 0.05.

| Gene Ontology (GO) | GO Term | Description | *p*-value | *q*-value |
|---|---|---|---|---|
| **Biological Process** | **GO:0043252** | **Sodium-independent organic anion transport** | **< 0.001** | **0.002** |
| | GO:0015711 | Organic anion transport | < 0.001 | 1.00 |
| **Molecular Function** | **GO:0015347** | **Sodium-independent organic anion transmembrane transporter activity** | **< 0.001** | **< 0.001** |
| **Cellular Component** | - | - | - | - |

**Table S11:** Gene Ontology (GO) enrichment for female relative fitness in the complex mating environment. GO categories enriched for genes annotated within 5kb of the leading haplotype block in each statistically significant genomic cluster (see Methods).

| Gene Ontology (GO) | GO Term | Description | *p*-value | *q*-value |
|---|---|---|---|---|
| **Biological Process** | - | - | - | - |
| **Molecular Function** | GO:0004180 | Carboxypeptidase activity | < 0.001 | 1.00 |
| **Cellular Component** | - | - | - | - |

**Table S12:** Gene Ontology (GO) enrichment for male relative fitness in the complex mating environment. GO categories enriched for genes annotated within 5kb of the leading haplotype block in each statistically significant genomic cluster (see Methods).

| Gene Ontology (GO) | GO Term | Description | *p*-value | *q*-value |
|---|---|---|---|---|
| **Biological Process** | - | - | - | - |
| **Molecular Function** | - | - | - | - |
| **Cellular Component** | GO:0043226 | Organelle | < 0.001 | 0.889 |
| | GO:0043229 | Intracellular | < 0.001 | 0.644 |
